# RHOV is a Detachment-Responsive Rho GTPase Necessary for Ovarian Cancer Peritoneal Metastasis

**DOI:** 10.1101/2025.08.12.669944

**Authors:** Amal T. Elhaw, Priscilla W. Tang, Ya-Yun Cheng, Shriya Kamlapurkar, Zaineb Javed, Sarah Al-Saad, Sierra R. White, Ahmed Emam Abdelnaby, Hannah Khan, Alex Seok Choi, Aidan R. Cole, Yeon-Soo Kim, Huda I. Atiya, Mohamed Trebak, Ioannis Zervantonakis, Ronald J. Buckanovich, Katherine M. Aird, Lan G. Coffman, Karthikeyan Mythreye, Nadine Hempel

## Abstract

All ovarian cancer subtypes spread via transcoelomic metastasis, where cells disseminate into the peritoneal fluid, resist anoikis, and form multicellular aggregates that invade the peritoneum. This represents the main driver of morbidity and mortality for peritoneal cancer patients. Mechanisms necessary for cancer cells to survive matrix detachment and initiate transcoelomic metastasis remain poorly defined. To address this, we identified a conserved detachment-sensitive gene signature activated shortly after matrix-detachment across multiple ascites-derived cancer cell lines. RHOV, an atypical, constitutively active and understudied member of the Rho GTPase family, emerged as a top upregulated transcript, which was confirmed in patient ascites-derived tumor cells. Functionally, loss of RHOV impairs anoikis resistance, multicellular aggregate integrity, migration and invasion, and completely abolishes transcoelomic tumor progression *in vivo*. RHOV enhances c-Jun signaling and cytoskeletal remodeling, which is dependent on both RHOV GTP-binding and membrane localization. These findings define RHOV as a novel detachment-sensitive Rho GTPase and establish RHOV as a critical regulator of peritoneal metastasis for the first time.

## Introduction

Epithelial ovarian cancer is the second most lethal gynecologic malignancy and the sixth leading cause of cancer-related deaths among women in the United States (1). This high mortality is largely attributed to the fact that over 70% of patients are diagnosed at an advanced, metastatic stage, where the five-year survival rate falls to approximately 30%. Regardless of histologic subtype, advanced stage patients present with widespread peritoneal dissemination and the presence of malignant ascites - features that reduce the effectiveness of standard cytoreductive surgery and chemotherapy (2–4). Given that metastasis accounts for the majority of OVCA-related deaths, there is an urgent need to elucidate the mechanisms driving peritoneal dissemination and to identify unique biological vulnerabilities that can be leveraged for future development of novel therapeutic strategies targeting metastatic disease.

Epithelial ovarian cancers, including high-grade and low grade serous, endometrioid, clear cell, and mucinous carcinomas, uniquely spread through transcoelomic metastasis, in which tumor cells directly disseminate from the primary tumor into the peritoneal cavity (2). Critical steps in transcoelomic metastasis include detachment of cells, resistance to anoikis (detachment-induced cell death) within the peritoneal fluid, and subsequent assembly into multicellular aggregates (MCAs) (5–7). These anoikis-resistant cells and aggregates function as metastasizing units, colonizing both neighboring and distant peritoneal sites and perpetuate recurrence via continuous shedding from both primary and secondary lesions. Indeed, the detection of anoikis-resistant cells or MCAs in patient ascites correlates with poor prognosis and chemoresistance (5, 8–11). Research on metastasizing ovarian cancer cells has largely focused on MCAs as metastasizing units; however, the immediate molecular adaptations triggered by matrix detachment, facilitating epithelial cell transition to an anchorage independent state and subsequent anoikis resistance and MCA formation, remain largely unexplored. This gap in knowledge limits our understanding of the early events that prime ovarian cancer cells for metastatic success. Our study takes a subtype-agnostic approach to investigate the initial adaptive responses that occur immediately after matrix detachment of metastasizing ovarian cancer cells, hypothesizing that these immediate early changes enhance the acclimatization of tumor cells as they transition from an adherent to a suspension state, ultimately enhancing metastatic fitness.

We employed both primary patient-derived ascites cells as well as three established ascites-derived ovarian cancer cell lines (OVCAR3, SKOV3, and OVCA433) to identify genes upregulated immediately following matrix detachment. Among these, the atypical Rho GTPase, RHOV, emerged as a consistent top hit across all models and was notably the sole Rho GTPase represented in this signature. Rho GTPases orchestrate cellular morphogenesis by translating extracellular mechanical cues into intracellular biochemical signals, an essential process during cellular state transitions (12–15). While the role of Rho GTPases has been described in the context of migrating and metastasizing cells with an emphasis on the most well characterized members (RHOA, RAC1, and CDC42) (14); little is known about the involvement of Rho GTPases in the initial steps following matrix detachment in general and specifically in the context of transcoelomic metastasis.

RHOV is an atypical Rho GTPase characterized by its rapid GTPase cycle that is hypothesized to render it almost constitutively active once expressed (16–18). RHOV has been studied primarily in developmental processes, such as Xenopus neural crest development, a phenomenon analogous to epithelial to mesenchymal transition (EMT), and zebrafish epiboly where RHOV regulates migration, adhesion, and differentiation (19–21). More recently, RHOV was found to be implicated in promoting EMT and metastatic progression in lung and breast cancer models (22–25). Despite these emerging findings, RHOV remains one of the least characterized Rho GTPases in tumor cells, and the functional relevance of RHOV in ovarian cancer remains entirely unexplored. In this study, we show for the first time that RHOV expression is rapidly induced in response to detachment, that RHOV is essential for anoikis-resistance, MCA compaction, migration and invasion of ovarian cancer cells and, most notably, is necessary for omental colonization *in vivo*, establishing RHOV as a critical mediator of transcoelomic metastasis. This work defines RHOV as a novel detachment-sensitive Rho GTPase and establishes RHOV as a critical regulator of ovarian cancer peritoneal metastasis, offering new insights into the molecular underpinnings of transcoelomic spread.

## Results

### A Conserved Detachment-Sensitive Transcriptome Highlights RHOV as a Unique Immediate Response Gene in Disseminating Ovarian Cancer Cells

Our previous work demonstrated that ovarian cancer multicellular aggregates (MCAs) cultured under anchorage independent (a-i) conditions regulate critical survival genes and stress adaptations essential for intraperitoneal metastasis (26, 27). Building on these findings, we sought to define the immediate early transcriptomic adaptations acquired by metastasizing cells following matrix detachment and preceding MCA formation. To this end, we performed high-depth RNA-sequencing of ovarian cancer cells at 2 hrs post-detachment, an interval selected to capture the nascent detachment-sensitive transcriptome of disseminating cells before secondary stress responses emerge. Based on the common nature of transcoelomic spread across serous epithelial ovarian cancer subtypes we employed ascites-derived NIH:OVCAR3 (TP53-mut-R248Q, CCNE1 amplified), OVCA433 (TP53-WT) and SKOV3 (TP53del-pS90PfsTer33) cell lines (28–30). Cells were profiled under three conditions: (i) attached monolayer culture (adherent state), (ii) single cell anchorage-independent state (2hr a-i), and (iii) MCA formation state (24 hrs a-i; Fig. 1A). Focusing on differential gene expression changes (log₂ fold-change >1, adjusted P <0.05) in response at 2hrs a-i revealed a conserved, detachment-sensitive gene signature shared across all three cell lines (Fig. 1A, B).

**Figure 1.**
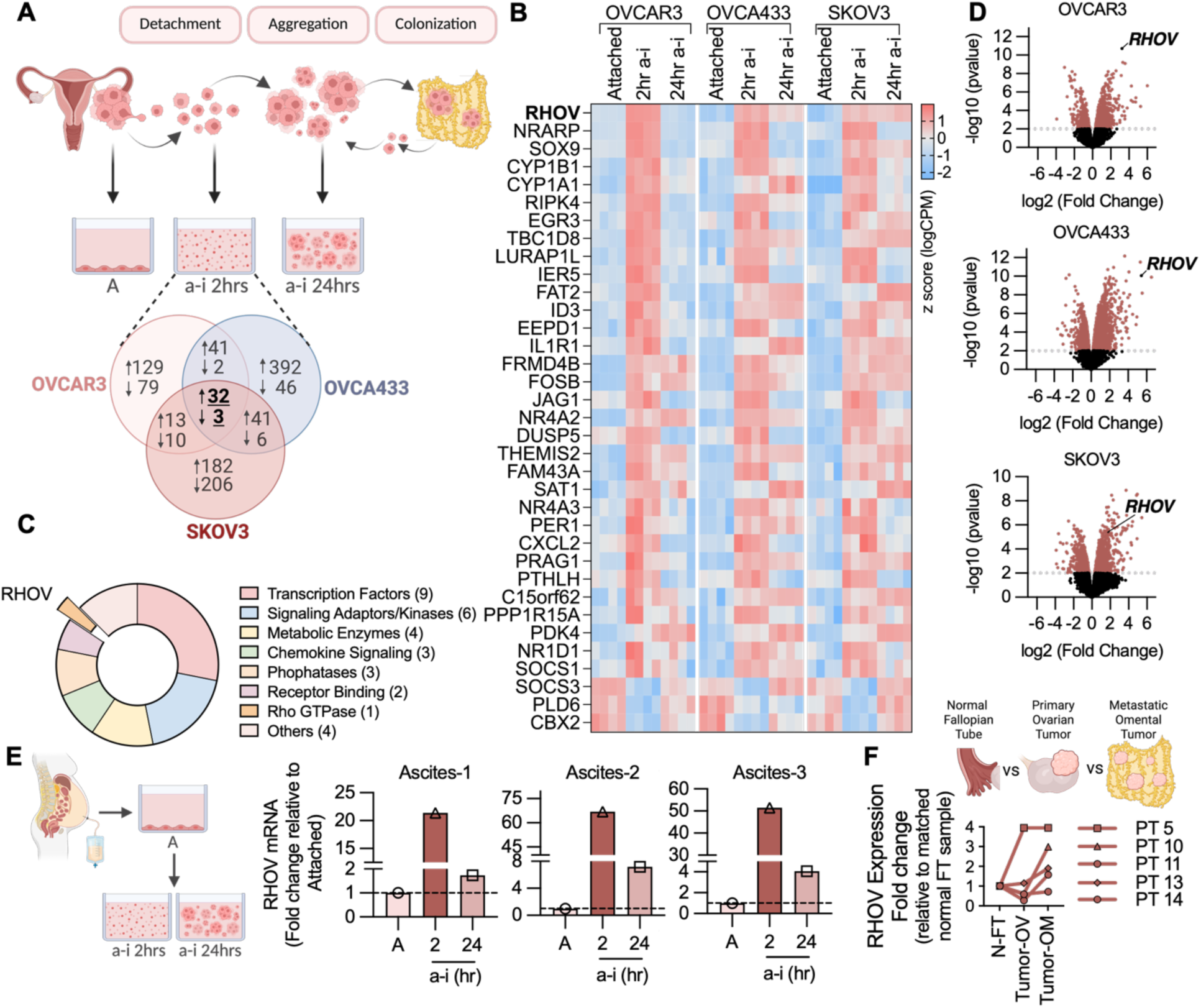
Detection of a detachment-sensitive transcriptomic signature in ovarian cancer cells led by the atypical Rho GTPase, RHOV, that is enriched in metastatic ovarian cancer tumors. **A,** Schematic overview of transcoelomic metastasis in ovarian cancer. Ascites-derived ovarian cancer cell lines (OVCAR3, OVCA433 and SKOV3 cells) were grown in tissue-culture treated dishes to mimic adherent (A) tumor cells. Cells were transitioned into anchorage independent condition (a-i) using ultra-low attachment plates for 2 hrs (a-i 2hrs), where they reside as single cells in suspension to mimic early disseminating cells, or for 24 hrs (a-i 24hrs), where cells form multicellular aggregates (MCAs). Venn diagram shows overlap of differentially expressed genes (n=4, log₂FC > 1, adjusted P < 0.05) across all three cell lines at 2hrs a-i. **B,** Heat map representing individual genes comprising the detachment-induced signature identified in (panel A) ranked based on significance in OVCAR3 cell line. **C,** Functional annotation of the conserved upregulated detachment-sensitive gene set highlighting RHOV as the sole Rho GTPase within the signature. **D,** Volcano plots of differentially expressed genes at 2 hrs in a-i showing that RHOV ranks among the top-most upregulated genes in all cell lines. **E,** RHOV is upregulated in ascites-derived tumor cells isolated from three high grade serous ovarian cancer patients when cultured under a-i conditions. **F,** RHOV is enriched in metastatic lesions in 4/5 high grade serous ovarian cancer patients compared to matched primary tumors and normal tissue (N-FT=Normal Fallopian Tube, Tumor-OV=Tumor Ovary and Tumor-OM=Tumor Omentum).

Functional annotation of the detachment-responsive gene signature revealed enrichment across diverse biological processes, including transcriptional regulation, metabolic pathways, chemokine signaling, and receptor-mediated signaling, and included genes such as NRARP, FOSB, IER5, and EGR3, well-characterized immediate early response genes that have been implicated in early metastatic dissemination in a hematogenous breast cancer model (31) (Fig. 1C; Fig. S1A), Within this signature, *RHOV* stood out as the sole Rho GTPase, ranking as the most significantly upregulated gene in OVCAR3 cells (Fig. 1D). Given the central role of Rho GTPases in translating extracellular biophysical cues into intracellular signaling, this selective upregulation of *RHOV* upon matrix detachment pointed to a potentially unique role for RHOV in ovarian cancer metastasis, which has not been previously explored. To test whether *RHOV* induction merely reflected a global activation of the Rho GTPase family, we examined expression changes for all 20 canonical Rho GTPases in our RNA-seq datasets. Strikingly, none displayed consistent and robust increased in transcript expression during early a-i compared to RHOV, including *RHOU*, the closest sequence homolog to *RHOV* (Fig. S2).. This demonstrates that RHOV’s induction in response to detachment is specific and non-redundant among Rho GTPases. Time-course analysis of *RHOV* transcript levels following culture in a-i demonstrated that *RHOV* transcripts were robustly elevated as early as 30 minutes post-detachment and reached a peak at around 2 hrs across all cell lines tested (Fig. S3A-C). The magnitude of induction was inversely correlated with basal RHOV expression (Fig. S3D), and RHOV expression was induced regardless of the method used for cellular detachment, including enzymatic and non-enzymatic dissociation, underscoring its generality as a detachment-triggered transcriptional response (Fig. S3E). Actinomycin-D completely abrogated the observed *RHOV* induction in early a-i (Fig. S3F), indicating that the detachment-induced surge in *RHOV* reflects *de novo* transcription rather than mRNA stabilization. Due to the understudied nature of RHOV, only a limited number of antibodies targeting RHOV are commercially available, all of which we found to exhibited poor signal-to-noise ratios and failed to reliably detect endogenous protein. However, polysome profiling revealed that *RHOV* transcripts rapidly redistributed into the heavy polysome fraction shortly after detachment indicating that RHOV transcripts are primed to be translated (Fig. S3G&H). Notably, RHOV is an atypical Rho GTPase distinguished by an accelerated GTPase cycle that is hypothesized to render RHOV constitutively active upon expression, suggesting that RHOV is regulated primarily at the level of transcription (16–18). Collectively, the above findings show that *RHOV de novo* transcription is rapidly deployed upon detachment and may represent an important regulatory mechanism of this atypical GTPase in cancer cells.

To validate the clinical relevance of the above findings, we isolated ascites-derived tumor cells from patients diagnosed with metastatic high grade serous ovarian cancer. Closely recapitulating the rapid upregulation observed in our cell line models, these primary tumor cells exhibited a marked increase in *RHOV* mRNA within 2 hours following detachment (Fig. 1E). We next assessed RHOV’s expression in matched clinical specimens from patients diagnosed with metastatic disease. In four of five matched sets, RHOV was significantly enriched in metastatic nodules compared to both the primary tumor and non-malignant control fallopian tube tissue (Fig. 1F), suggesting that cells with elevated RHOV during initial detachment gain a selective advantage in successful metastatic colonization to the omentum.

### Genetic Ablation of RHOV Impairs Transcoelomic Dissemination and Peritoneal Colonization of Ovarian Cancer Cells *in vivo*

To elucidate the functional role of RHOV in disseminating ovarian cancer cells, we established isogenic RHOV-knockout (KO) derivatives from the three ascites-derived ovarian cancer cell lines used in our initial screen. Utilizing a multiplex sgRNA CRISPR–Cas9 strategy, we targeted *RHOV* exon 1, inclusive of its translation start site (Fig. S4A). Genomic PCR using primers flanking the targeted locus identified clones with complete RHOV deletion in OVCA433 and SKOV3 cells, whereas OVCAR3 cells, displayed a partial deletion (Fig. S4B, C). RT-PCR confirmed the loss of detectable RHOV transcripts in both attached and a-i conditions in OVCA433-KO and SKOV3-KO clones and partial KO in OVCAR3 cells (Fig. S4D).

We first investigated whether RHOV is critical for transcoelomic ovarian cancer metastasis using two complementary *in vivo* models. To assess RHOV’s role in early metastatic dissemination from the ovarian niche an orthotopic intra-bursal xenograft model was employed using parental wild type (WT) and partial RHOV-KO OVCAR3 cells (Fig. 2A). Following ovarian tumor establishment, OVCAR3-WT cells efficiently disseminated throughout the peritoneum. Conversely, OVCAR3-RHOV-KO tumors remained confined to the primary ovarian site for the study duration (Fig. 2B, C), with complete absence of peritoneal lesion in the OVCAR3 RHOV-KO group (Fig. 2D), indicating that RHOV is a critical mediator of metastatic progression from the primary ovarian site. Ex-vivo analysis showed that ovarian bioluminescence was reduced in RHOV-KO versus WT tumors, and omental metastasis was completely abrogated by RHOV-KO (Fig. 2E, F). Comparing *in vitro* growth rates of WT and RHOV-KO cells across the three ovarian cancer cell lines demonstrated only modest, cell line–dependent reductions (Fig. S5), suggesting that inherent changes in proliferation in response to RHOV KO do not contribute to the dramatic decreases in metastatic tumor burden observed *in vivo*.

**Figure 2.**
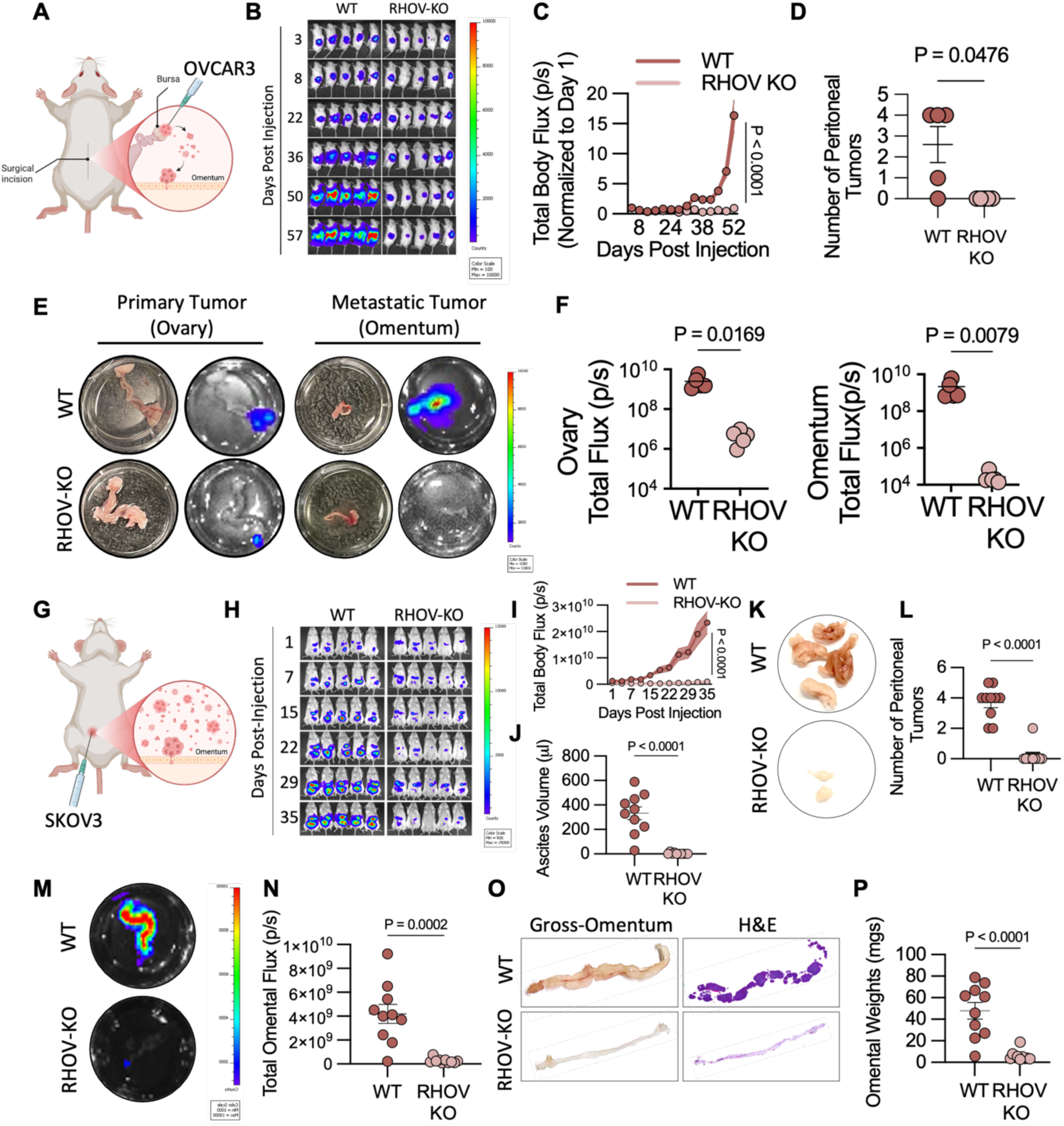
Genetic deletion of RHOV impairs transcoelomic dissemination and peritoneal colonization of ovarian cancer cells. **A,** Schematic of orthotopic intra-bursal xenograft model. Luciferase-labeled OVCAR3 cells were injected into the left ovary-bursa of NSG mice. **B,** Representative bioluminescence images of mice post-intra-bursal tumor cells injection. OVCAR3 WT tumors disseminate widely where RHOV-KO tumors remain confined to the ovary. **C,** Quantification of total peritoneal bioluminescence from (B), (n=5 mice/group, two way ANOVA, P < 0.0001). **D,** Quantification of peritoneal tumors in OVCAR3-WT vs RHOV-KO mice (n=5; Unpaired t-test P value shown). **E,** Representative gross (left) and bioluminescence (right) images of post-necropsy ovary and omental tumors. Mice injected with OVCAR3 RHOV-KO cells show no signal at omental metastatic sites. **F,** Quantification of ovary and omental bioluminescence signal from (E), demonstrating reduced primary tumor burden from RHOV-KO OVCAR3 cells as well as a complete absence of metastasis to the omentum (n=5; Ovary: Unpaired t-test P-value shown, Omentum: Mann Whitney test P-value shown). **G,** SKOV3 cells were injected directly into the peritoneal cavity in an a-i state to mimic transcoelomic metastasis of pre-disseminated cells. **H,** Representative in vivo bioluminescence images of NSG mice following intraperitoneal injection with luciferase-labeled SKOV3 WT or RHOV-KO cells. **I,** Quantification of total photon flux from whole-body imaging in (H), showing significantly reduced metastatic burden in RHOV-KO–tumor bearing mice (n=10 mice; two way ANOVA P < 0.0001). **J,** Volume of ascites fluid collected from SKOV3 WT and RHOV-KO group (n=10, Unpaired t-test, P value shown). **K, L,** Representative images and quantification of number of peritoneal tumor nodules in SKOV3 WT and RHOV-KO injected mice showing reduced size and incidence of peritoneal metastases in RHOV-KO group (n=10, Unpaired t-test P value shown). **M,** Representative ex vivo bioluminescence images of excised omentum from mice injected with SKOV3-WT and KO cells showing reduced flux in the RHOV-KO mice omentum. **N,** Quantification of omental bioluminescence intensity from (M), (n=10, Unpaired t-test P value shown). **O,** Representative gross images and H&E staining of omentum from SKOV3 WT and RHOV-KO injected mice. **P,** Quantification of omental weight from SKOV3 WT and RHOV-KO injected mice (n=10, Unpaired t-test P value shown).

In addition, an intraperitoneal (IP) xenograft metastasis model using SKOV3 WT and RHOV-KO cells was employed, where cells were directly introduced into the peritoneal cavity in an anchorage-independent state within two hours of their detachment (Fig. 2G). This model specifically assesses the competence of disseminated cells forced into suspension to resist anoikis and colonize the peritoneal surfaces following injection. NSG mice injected with SKOV3 RHOV-KO cells exhibited significantly reduced bioluminescent signal intensity and spatial metastatic distribution compared to their WT counterparts (Fig. 2H, I). Consistent with diminished peritoneal colonization, RHOV-KO tumor bearing mice also accumulated substantially less ascites (Fig. 2J). Moreover, both the size (Fig. 2K) and the frequency (Fig. 2L) of peritoneal tumor nodules were significantly lower in RHOV-KO xenografts. *Ex vivo* imaging (Fig. 2M, N) and histological evaluation (Fig. 2O, P) of excised omentum further confirmed negligible tumor cell colonization by RHOV-KO tumor cells. These data underscore that RHOV is essential for the metastatic colonization of suspended ovarian cancer cells within the peritoneal cavity. These in vivo findings suggest that RHOV’s activity during early anchorage independence may have lasting effects on metastatic progression, supporting a potential role for RHOV in facilitating the metastatic potential of ovarian cancer cells.

### Detachment-Induced RHOV Expression Mediates Anoikis Resistance

Having established RHOV’s critical role in peritoneal metastasis *in vivo*, we next sought to delineate the functional contributions of RHOV throughout distinct stages of the ovarian cancer metastatic cascade. Given RHOV’s robust induction following detachment, we investigated the role of RHOV in mediating single cell survival in a-i, a fundamental prerequisite for transcoelomic metastasis (5). OVCA433 cells were cultured in a-i as dispersed single entities or small clusters, using flat bottom ultra-low attachment (ULA) plates, and cell viability assessed by live/dead staining with Calcein AM/Ethidium Homodimer, respectively (Fig. 3A). While RHOV-KO did not influence cell viability in adherent conditions (Fig. S5B), RHOV-KO significantly increased susceptibility to anoikis, with a continuous rise in cell death up to 72 hrs in a-i conditions (Fig. 3B-D). These results suggest that RHOV plays an immediate protective effect during early detachment that is necessary to prime the cells for prolonged cell viability under anchorage-independent conditions. Additionally, clonogenic survival assays further confirmed that RHOV-KO cells had markedly reduced colony-forming capacity when seeded as dispersed single cells, reinforcing RHOV’s key role in mediating single cell survival post-detachment (Fig. S6A).

**Figure 3.**
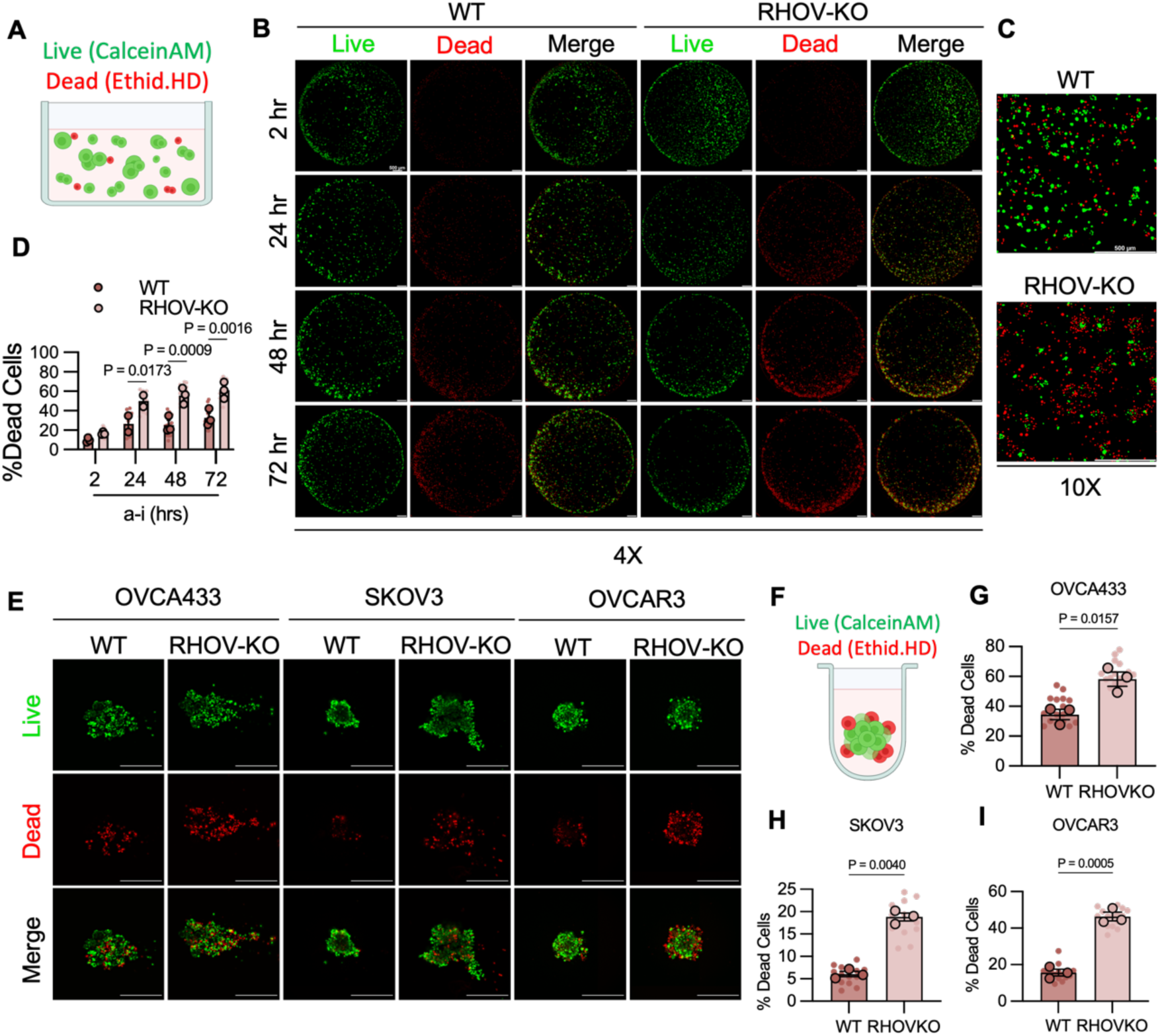
Detachment-induced RHOV expression promotes anoikis resistance in single ovarian cancer cells and multicellular aggregates. **A,** Live/dead staining of OVCA433 cells cultured under anchorage-independent (a-i) conditions in flat bottom ultra-low attachment plates (ULA) in the absence of serum keeps cells dispersed as single cells and small clusters in suspension, Calcein-AM (green) marks live cells; ethidium homodimer (red) marks dead cells. **B,** Representative fluorescent images of Live/Dead WT and RHOV-KO OVCA433 cells over time in anchorage-independent conditions in flat bottom ULA. Images were taken at 4x magnification and stitched (2×2 stich) to show whole well. Scale Bar=500uM. **C,** 10x magnification of Live/Dead cells from OVCA433 WT and RHOV-KO cells in flat bottom ULA from 72Hrs in a-i, scale bar=500uM. **D,** RHOV-KO cells show significantly increased single cell anoikis susceptibility. Time-course quantification of cumulative cell death under a-i conditions in OVCA433 WT and RHOV-KO cells in flat bottom ULA model (n=3, two way ANOVA P <0.0001, Sidak’s multiple comparisons post hoc test P value shown). **E,** Representative images of multicellular aggregates (MCAs) formed by OVCA433, SKOV3, and OVCAR3 WT and RHOV-KO cells after 72 hrs in round-bottom ULA plates imaged at 10x magnification, Scale Bar=500uM. **F,** Cells cultured under anchorage-independent (a-i) conditions in round bottom ultra-low attachment plates (ULA) in the presence of serum allowing cells to cluster into multicellular aggregates in suspension, Calcein-AM (green) marks live cells and ethidium homodimer (red) marks dead cells. **G-I,** Quantification of Live/dead staining of MCAs at 72 hrs in a-i reveals significantly increased cell death in RHOV-KO cell aggregates in OVCA433 (G), SKOV3, (H), OVCAR3 (I) (n = 3, Unpaired t-test P value shown).

Subsequent to initial detachment and single cell anoikis resistance, ovarian cancer cells typically aggregate into multicellular aggregates (MCAs) which promotes collective anoikis resistance and metastatic efficiency (10, 32, 33). To test the effect of RHOV-KO on MCA anoikis resistance, MCAs were generated by culturing cells in round-bottom ULA plates to promote aggregation (Fig. 3F). Consistent with single-cell observations, live/dead staining of MCAs revealed significantly increased sensitivity to anoikis in RHOV-KO aggregates compared to their wild-type counterparts across all tested cell lines (Fig. 3E-I). siRNA-mediated RHOV depletion in OVCA433 and SKOV3 similarly increased cell death and significantly elevated caspase activity under a-i conditions, further validating RHOV’s crucial role in mediating anoikis resistance following matrix-detachment (Fig. S7A, B). Together, these findings illustrate that RHOV induction during early detachment is critical for both immediate single-cell a-i survival and continued viability of MCAs (Fig. 4H; i).

**Figure 4.**
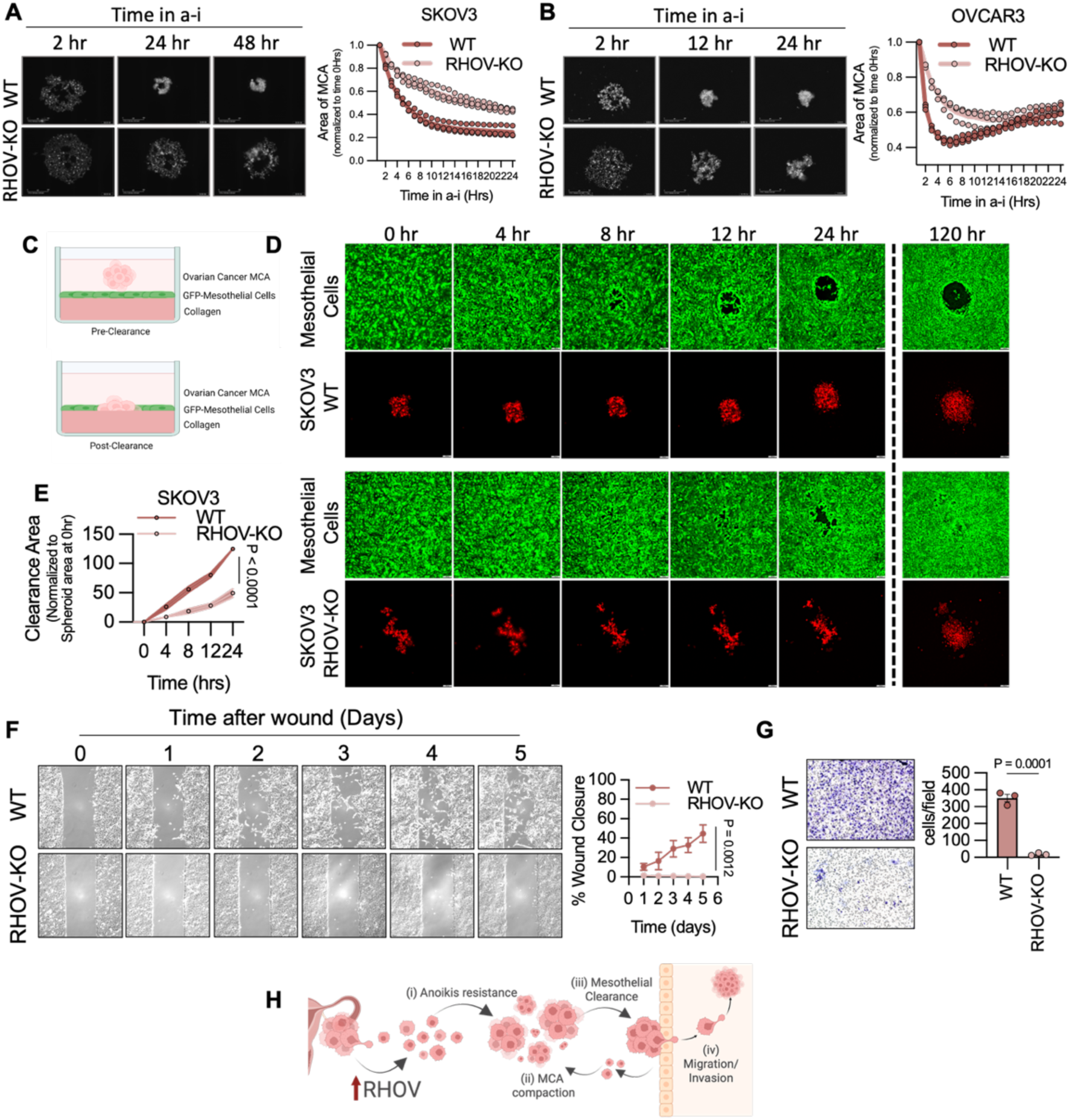
RHOV promotes MCA compaction, mesothelial clearance, and invasive behavior in ovarian cancer cells. **A,** MCA spheroid compaction assay in SKOV3 WT and RHOV-KO cells, showing reduced compaction in KO cells (n=3, two way ANOVA, P < 0.0001). **B,** MCA spheroid compaction assay in OVCAR3 WT and RHOV-KO cells, showing reduced compaction in KO cells (n=3, two way ANOVA, P < 0.0001). **C,** Schematic of the mesothelial clearance assay. RFP-labeled ovarian cancer spheroids were co-cultured atop a GFP-expressing mesothelial monolayer on collagen to mimic peritoneal colonization. **D,** Representative images of mesothelial clearance at indicated timepoints. RHOV-KO spheroids show significantly impaired clearance compared to WT cells, failing to fully displace the mesothelium or engage with the ECM. Images acquired at 10x magnification, Scale bar= 100uM. **E,** Quantification of mesothelial clearance area over time. RHOV-KO spheroids show progressive failure to expand and maintain cleared zones (n=3; two way ANOVA P value shown). **F,** Representative images and quantification of wound healing assay in SKOV3 WT and RHOV-KO cells showing loss of migratory capacity in RHOV-KO cells, Images acquired at 10x magnification, scale bar=100uM (n=3; two way ANOVA P value shown). **G,** Transwell invasion assay demonstrates severely reduced Matrigel invasion in RHOV-KO SKOV3 cells (n=3, unpaired t-test P value shown). **H,** Summary schematic illustrating RHOV’s role across sequential steps of metastatic progression.

### RHOV Is Required for Structural Integrity and Invasive Behavior of Ovarian Cancer Cell MCAs

MCAs play a pivotal role in invading the peritoneum and colonizing peritoneal organs, processes shown to be influenced by the physical properties of the spheroids, including their compaction (34, 35). While analyzing spheroid viability, we noted that RHOV-KO MCAs often appeared morphologically loose and poorly compacted (Fig. 3E). To assess RHOV’s role in regulating spheroid architecture, a spheroid compaction assay was carried out to measure the reduction in spheroid diameter over the first 24 hrs of anchorage-independent culture. In both SKOV3 and OVCAR3 lines, which form compact spheroids in a-i (Fig. 3E), RHOV knockout significantly impaired compaction compared to WT controls (Fig. 4A, B). These findings suggest that RHOV’s role in early a-i facilitates subsequent MCA compaction (Fig. 4H; ii).

We next investigated whether RHOV’s effect on MCA compaction functionally impairs their metastatic potential. A mesothelial clearance assay was used, in which pre-formed, RFP-labeled ovarian cancer MCAs were co-cultured atop a confluent monolayer of GFP-expressing mesothelial cells on a collagen matrix, recapitulating the *in vivo* microenvironment of the omentum, the principal site of ovarian cancer metastasis (Fig. 4C). This assay models mesothelial cell displacement by cancer cells and their subsequent engagement with the submesothelial extracellular matrix required for implantation (36–39). Using SKOV3 cells, the most invasive line in our panel, we found that RHOV-depleted MCAs were markedly impaired in their ability to clear the mesothelium (Fig. 4D, E). Notably, after prolonged co-culture, RHOV-KO spheroids failed to maintain mesothelial clearance, appearing to grow atop the intact mesothelial monolayer rather than successfully displacing it or competing for binding to the collagen substrate below. This highlights a key deficiency in the metastatic competence of RHOV-deficient MCAs (Fig. 4H; iii).

To further dissect the downstream cellular behaviors influenced by RHOV related to tissue intravasation, we examined its role in the regulation of migratory and invasive phenotypes. RHOV-KO in SKOV3 cells, which exhibit the highest RHOV expression in attached conditions (Fig. S3D) displayed a complete inability to migrate into the wound gap in a scratch assay (Fig. 4F), which was similarly observed by siRNA-mediated RHOV knockdown (Fig. S7C). Complementary transwell invasion assays revealed that RHOV-KO cells also exhibited severely reduced invasion through Matrigel (Fig. 4G). These findings uncover a novel role of RHOV in mediating ovarian cancer migration and invasion, processes crucial during later stages of metastatic colonization (Fig. 4H; iv). Taken together, we show that RHOV is a pivotal regulator of sequential metastatic events that follow the detachment of ovarian cancer cells, including anchorage-independent survival, spheroid compaction, mesothelial clearance, migration and invasion (Fig. 4H).

### RHOV Regulates Pro-Metastatic Transcriptional Programs in Early Anchorage-Independence

To dissect the mechanistic underpinnings of RHOV-mediated ovarian cancer metastasis, we performed bulk RNA sequencing following RHOV-KO. OVCA433, which demonstrate robust induction of RHOV expression upon detachment, were cultured under three conditions to capture RHOV-dependent transcriptomic changes under adherent conditions, following immediate detachment (2 hrs a-i) and in MCAs (24 hrs a-i). We observed distinct roles for RHOV in governing transcriptional changes under these different conditions. In adherent cells, Gene Ontology (GO) enrichment analysis revealed that RHOV loss was associated with downregulation of cytokine-mediated signaling, including interleukin and interferon responses (Fig. 5A, Fig. S8A). However, in early anchorage-independent conditions (2 hrs a-i), RHOV-KO cells displayed robust suppression of pro-metastatic pathways including cell-cell junction organization, positive regulation of cell adhesion, cell migration, and negative regulation of apoptosis, (Fig. 5B). Downregulation of these pathways correlates with the observed phenotypes in RHOV-KO cells, such as diminished survival in anchorage-independent conditions, impaired spheroid compaction, and reduced migratory capacity. Genes driving the topmost downregulated pathway included *MYLK*, a key regulator of actomyosin contractility, as well as *SERPINE1, F2RL1, and F2R*, all of which are known to govern migratory signaling in cancer cells (Fig. S8B) (40–43).

**Figure 5.**
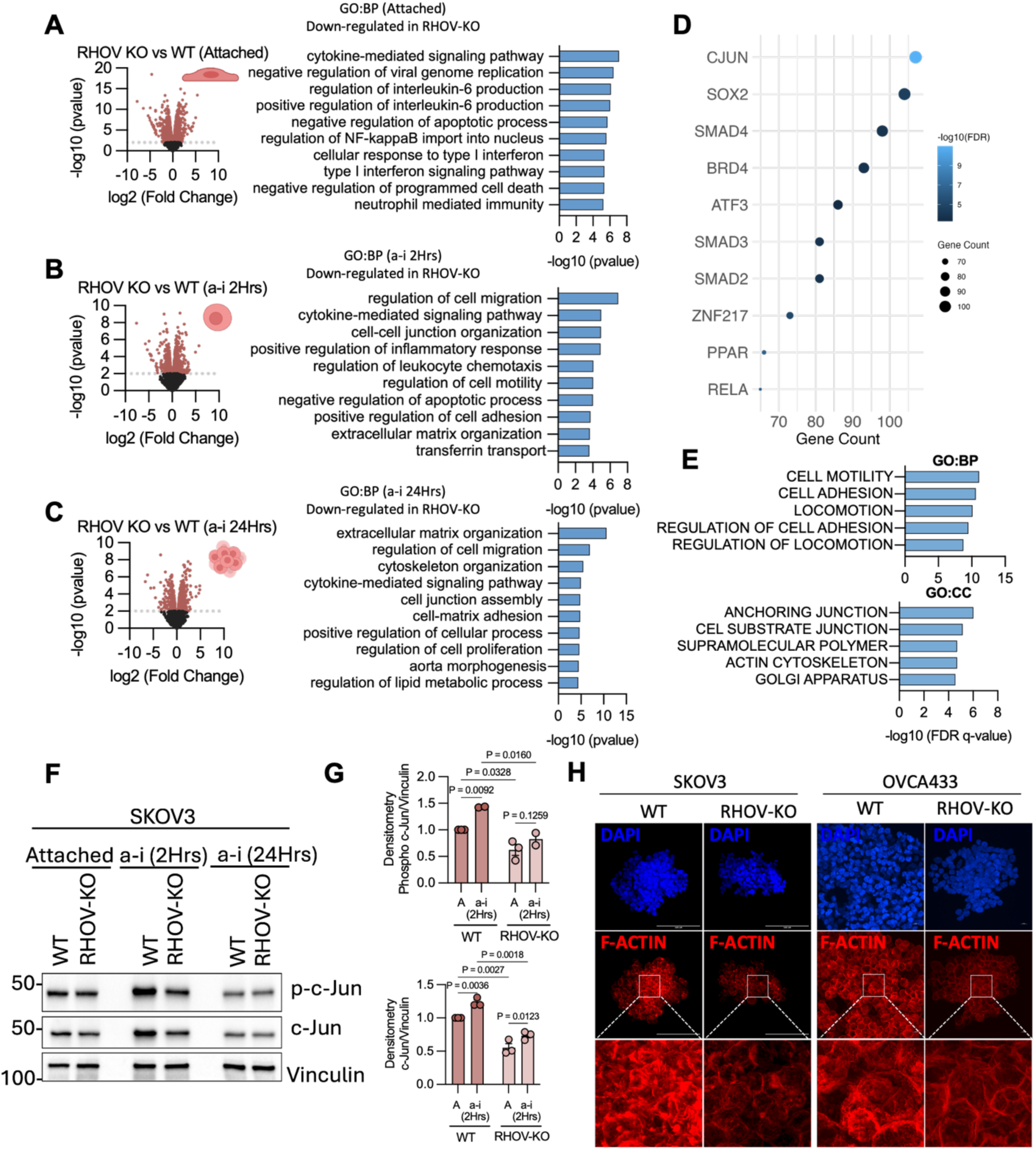
RHOV regulates c-Jun dependent cytoskeletal remodeling in ovarian cancer cells. **A–C,** Bulk RNA-sequencing of OVCA433 WT and RHOV-KO cells cultured under three conditions: adherent (A), early anchorage-independent (2 hrs; B), and late anchorage-independent (24 hrs; C). Left panels: volcano plots showing distribution of differentially expressed genes between WT and KO cells. Right panels: Gene Ontology (GO) Biological Pathways enrichment analysis of significantly downregulated pathways in RHOV-KO cells in each condition (n=3). **D,** Transcription factor enrichment analysis (TFEA) of differentially expressed genes at the 2 hrs a-i timepoint identifies c-Jun as the top predicted downregulated transcriptional regulator in RHOV-KO cells. **E,** GO enrichment of RHOV dependent c-Jun–regulated genes shows their convergence on pathways related to actin cytoskeleton organization, anchoring junctions, and cell motility. **F-G**, Western blots representative image (F) and quantification (G) showing reduced phospho-c-Jun (Ser63) in RHOV-KO SKOV3 cells (n=3, two way ANOVA P values shown). **H,** Phalloidin staining shows reduced polymerized F-actin in RHOV-KO cells under early a-i (2hrs) conditions, indicating impaired actin cytoskeletal remodeling in both OVCA433 and SKOV3 RHOV-KO cells. Images acquired at 63x magnification. Scale bar = 100uM (SKOV3), 10uM (OVCA433).

At the 24-hrs a-i condition, where cells transition into MCAs, the RHOV-regulated transcriptome revealed downregulation of genes associated with extracellular matrix (ECM) organization, cytoskeletal organization, and cell-matrix adhesion (Fig. 5C). The leading-edge genes within the top pathway included several ECM structural components (*COL4A1, COL4A2*) and a broad repertoire of integrins (*ITGA1, ITGA3, ITGA5*, and *ITGA7*) all of which are critical for mediating spheroid compaction, binding to collagen underlying mesothelium, and subsequent invasion (Fig. S8C) (44–46). These transcriptional changes align closely with the phenotypic defects observed in RHOV-KO MCAs, including compromised spheroid survival, defective mesothelial clearance, and impaired adhesion to the mesothelial surface and subsequent invasiveness within the peritoneal cavity. Together, these results point to a temporally coordinated role for RHOV in regulating metastatic progression, where early transcriptional changes during 2hrs a-i influence key processes such as migration, adhesion, and apoptosis. These early adaptations may subsequently shape the invasive behavior and collagen-binding capacity observed in MCAs, suggesting that RHOV-driven programs during the initial detachment phase contribute to the establishment of long term pro-metastatic traits.

### RHOV Orchestrates c-Jun Signaling and Actin Cytoskeletal Remodeling in Disseminating Ovarian Cancer Cells

To uncover key effectors downstream of RHOV that mediate RHOV-dependent transcription and pro-metastatic function during early dissemination, we focused on the 2 hrs a-i timepoint, where RHOV expression peaks and RHOV-driven pro-metastatic transcriptional programs are first induced (Fig. 5B). Transcription factor enrichment analysis (TFEA) revealed c-Jun as the top-ranked transcription factor predicted to be functionally suppressed in RHOV-KO cells in early a-i (Fig. 5D). Gene ontology analysis of RHOV-dependent c-Jun–associated transcripts again revealed a strong enrichment in pathways regulating cell motility, anchoring junction assembly, and actin cytoskeleton organization (Fig. 5E). These findings position c-Jun as a potential effector of the RHOV pro-metastatic transcriptional response. This was confirmed by immunoblotting in both OVCA433 and SKOV3 cells, demonstrating increased c-Jun (S63) phosphorylation in early a-i and a significant reduction in phospho-c-Jun in RHOV-KO cells at 2 hrs post-detachment (Fig. 5F-G, Fig. S8D). Total c-Jun levels were also reduced, aligning with reports that phospho-c-Jun can enhance its own transcription through positive feedback mechanisms (47). The above data support RHOV’s role in activating c-Jun signaling (23, 48), but demonstrate for the first time that this RHOV-dependent signaling axis is functionally engaged in metastatic ovarian cancer cells following detachment. Given the convergence of RHOV and c-Jun on pathways governing cytoskeletal dynamics, we assessed polymerized F-actin assembly by phalloidin staining at the 2-hour a-i time point in both OVCA433 and SKOV3 cells. RHOV-KO cells exhibited markedly reduced polymerized F-actin intensity relative to their WT counterparts (Fig. 5H). These data show that RHOV facilitates actin remodeling following cancer cell detachment that may be necessary during early stages of dissemination.

### RHOV’s GTP-binding and Membrane Localization Are Required for Signaling and Pro-metastatic Function

To define the molecular features essential for RHOV’s function, we rescued RHOV-KO cells with wild-type (WT) RHOV and three targeted mutants: constitutively active (CA; GTP-bound; G40V); dominant-negative (DN; GDP-bound; S45N), and a membrane localization-deficient (ΔMEM) mutant (C234S), which harbors a cysteine-to-serine substitution in the conserved CFV motif that blocks palmitoylation and membrane targeting, while preserving GTPase cycling activity (Fig. 6A) (49). These constructs were stably expressed in RHOV-KO SKOV3 cells to evaluate their capacity to rescue key RHOV-dependent functions. Both wild-type RHOV (WT) and the constitutively active (CA; G40V) mutant restored resistance to detachment-induced cell death, indicating that RHOV GTP-binding is required for anchorage-independent survival. In contrast, neither the dominant-negative (DN; S45N) mutant nor the membrane localization–deficient (ΔMEM; C234S) mutant conferred any survival advantage under a-i conditions (Fig. 6B, C), suggesting that both GTP-binding and membrane targeting are essential for this function. A similar pattern was observed in wound healing assays: only RHOV-WT and CA restored migratory capacity in RHOV-KO cells, while DN and ΔMEM remained ineffective (Fig. 6D, E). Notably, the indistinguishable behavior of RHOV-WT and the GTP-locked CA mutant across both assays provides direct functional evidence that WT RHOV behaves as a constitutively active Rho GTPase under physiological conditions.

**Figure 6.**
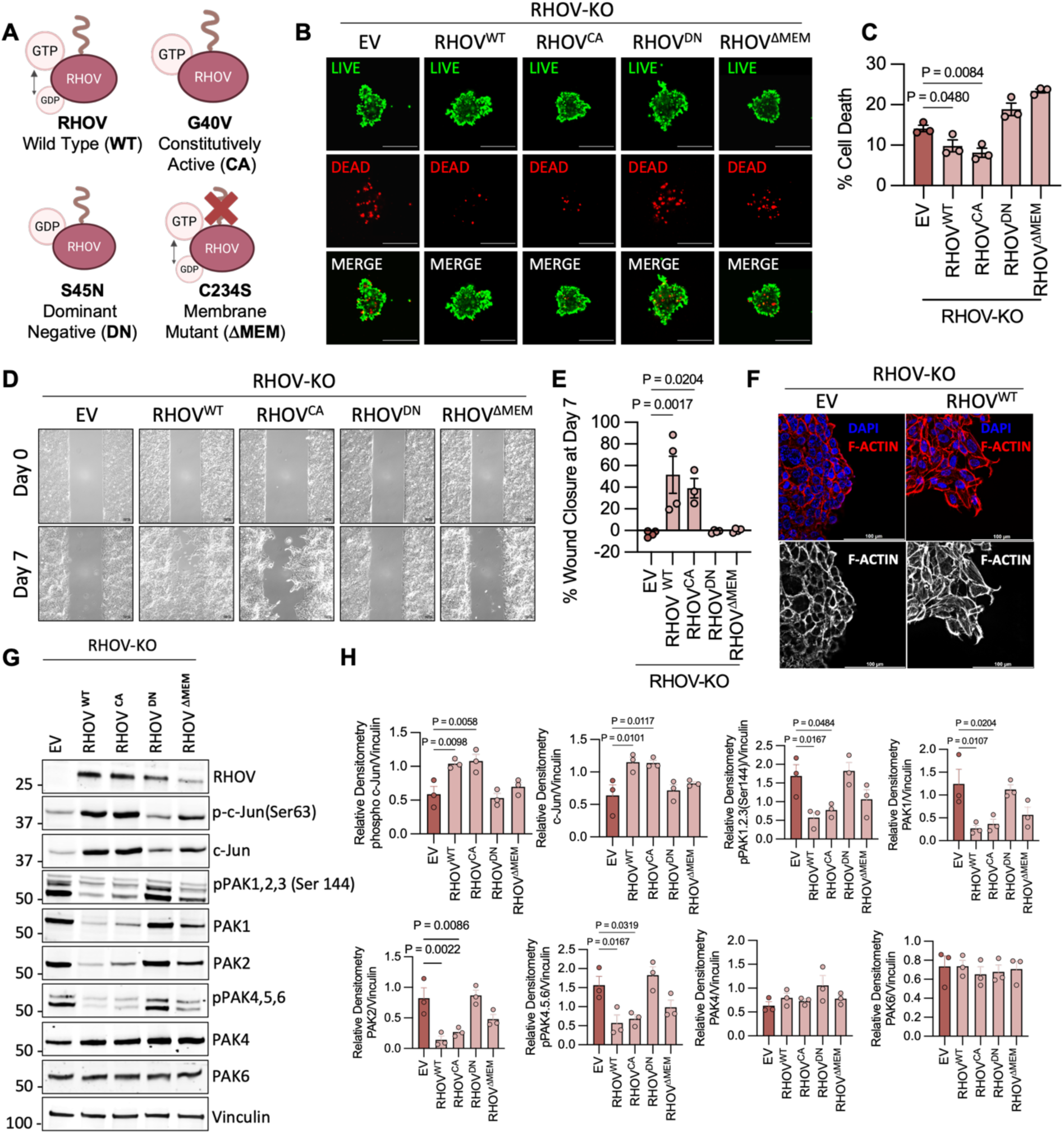
RHOV’s GTPase activity and membrane localization are required to restore cytoskeletal, transcriptional, and pro-metastatic functions in RHOV-deficient ovarian cancer cells. **A,** Schematic of RHOV constructs used in rescue experiments: wild-type RHOV (WT), GTP-locked constitutively active G40V mutant (CA), dominant-negative GDP-bound S45N mutant (DN), and palmitoylation-deficient membrane localization C234S mutant (**ϕλ**MEM). **B,** WT and CA RHOV but not DN or **ϕλ**MEM RHOV-mutants restore anoikis resistance in RHOV-KO SKOV3 aggregates after cells were cultured in anchorage independent (a-i) conditions for 72Hrs using ultra-low attachment plates. Representative images of live cells are labeled with Calcein-AM (green), and dead cells with Ethidium Homodimer (red). Images acquired at 10x magnification. Scale bar = 500uM. **C,** Quantification of live/dead percentage from (B), presented as percentage of dead cells relative to total (n=3; One-way ANOVA P <0.0001, Sidak’s post hoc test P value shown). **D,** Representative wound healing assay images at 0- and 7-days post-scratch in RHOV-KO SKOV3 cells expressing indicated RHOV rescue constructs or EV control. **E,** Quantification of % wound closure in (D) at day7; RHOV-WT and CA restore migratory capacity, while DN and **ϕλ**MEM do not (n=3; One-way ANOVA P=0.0009, Sidak’s post hoc test P value shown). **F,** Phalloidin staining of F-actin reveals restoration of actin architecture in RHOV-WT expressing SKOV3-RHOV-KO cells. Images acquired at 63x magnification. Scale bar = 100uM. **G, H,** Characterization of c-Jun/Pak pathway components in SKOV3 RHOV-KO+EV vs RHOV-rescue mutants. Representative Western blots are shown. Densitometry of protein bands from replicate experiments are quantified (n=3; One-way ANOVA, Sidak’s post hoc test P value shown).

We next asked whether the restoration of anoikis resistance and migratory capacity in RHOV-rescued cells is accompanied by re-establishment of actin architecture. Phalloidin staining revealed that expression of WT-RHOV in RHOV-KO cells reinstated actin polymerization, consistent with RHOV-dependent cytoskeletal remodeling (Fig. 6F). This was further accompanied by reactivation of c-Jun, a key driver of cytoskeletal transcriptional programs downstream of RHOV (Fig. 6G, H). As with earlier assays, RHOV-DN and ΔMEM did not rescue c-Jun phosphorylation unlike RHOV-WT and CA constructs. Given that RHOV-induced c-Jun activation likely involves upstream kinase signaling, and prior studies have implicated RHOV in regulating p21-activated kinases (PAKs) (16, 25, 50–52), we investigated how RHOV-rescue affects PAK phosphorylation. In RHOV-WT and CA expressing cells, we observed a striking reduction in phospho-PAK1/2/3 (Ser144), accompanied by a loss of total PAK1 and PAK2 protein levels (PAK3 was undetectable) (Fig. 6G–H). This aligns with previous reports suggesting that following PAK activation RHOV induces PAK1 autophosphorylation–dependent degradation (52) and further extends this RHOV-dependent regulatory mechanism to additional group I PAKs in ovarian cancer cells. To confirm that the observed reductions in PAK levels were due to rapid proteasomal degradation, we treated cells with the proteasome inhibitor MG132 (Fig. S9). This restored both phospho- and total PAK1/2/3 levels in WT and CA expressing cells, further supporting the activation-dependent degradation model (52). We also observed a reduction in phospho-PAK4/5/6 in RHOV-WT and CA cells, while total PAK4 and PAK6 protein levels remained unchanged (PAK5 was undetectable). This is intriguing as group II PAK activity is regulated by GTPases but their phosphorylation is typically GTPase–independent (53), and RHOV was previously shown to interact with PAK6, but not alter its phosphorylation (51).

The above data collectively demonstrate that RHOV’s ability to drive metastatic behavior requires both its GTP-bound state and proper membrane localization. When these features are intact, RHOV promotes anoikis resistance, migration, actin cytoskeletal remodeling, and signaling via PAK/c-Jun. Together, these results support a model in which RHOV operates as a constitutively active, membrane-associated Rho GTPase that is highly induced in response to detachment during peritoneal metastasis.

## Discussion

The majority of ovarian cancer patients are diagnosed at advanced stage, where metastasis is already established widely within the peritoneal cavity. These peritoneal tumors contribute to malignant ascites, a clinical feature that is associated with poor patient outcomes (4). As a result, attention has been directed toward understanding the adaptations acquired by metastasizing tumor cells within the peritoneal ascites, particularly those that enable multicellular aggregates (MCAs) to survive in suspension, persist, and colonize secondary sites (26, 27, 44, 54–56). While numerous studies have defined key stress response programs that sustain metastatic fitness within spheroids, the earliest cellular responses immediately following detachment from the extracellular matrix have remained largely unexamined. This critical transition period represents a poorly understood but biologically distinct phase of metastatic progression. Our findings underscore the importance of investigating these acute detachment-induced adaptations, which we show to play a foundational role in priming disseminating cells for subsequent anoikis resistance, invasion, and peritoneal colonization.

Our work indicates that early transcriptomic adaptations following detachment are not only distinct but also essential for downstream metastatic success. We identified a conserved, detachment-sensitive gene signature activated at 2 hrs in a-i culture conditions, and focused on RHOV, the sole Rho GTPase and most significantly upregulated gene in OVCAR3 cells. Importantly, RHOV induction was validated in primary tumor cells isolated from patient ascites, supporting the pathophysiological relevance of this response. The functional importance of this rapid transcriptional upregulation was immediately apparent by the observation that genetic ablation of RHOV profoundly impaired metastatic progression *in vivo*. These findings establish RHOV not merely as a marker of detachment-induced stress, but as a critical effector of early dissemination and peritoneal colonization, whose induction is essential for successful metastatic progression. We hypothesize that the detachment responsive gene expression signature reflects an acute stress response adopted by detaching ovarian cancer cells, as it includes a number of immediate early genes, including the transcription factors FOSB, IER5, and EGR3, which are known stress induced genes largely studied in neuronal plasticity and memory formation (57). Given the rapid and robust transcriptional regulation, RHOV shares these features with immediate early genes. Although less is known about their function in tumor cells, several of these immediate early genes were recently independently identified in an i*n vivo* model of early hematogenous metastatic spread in breast cancer (31). This further supports the potential importance of the detachment response signature identified here in the metastatic process and opens future studies investigating the role of these genes in peritoneal metastasis.

Unlike classical Rho GTPases, whose activity is dynamically modulated by Guanine nucleotide exchange factors (GEF) and GTPase-activating proteins (GAP), RHOV (also known as Chp (Cdc42 homologous protein)/Wrch-2 protein) is predicted to exist in a constitutively GTP-bound state and bypass canonical cycling-based control (16–18). Instead, its activity is thought to be governed through tightly regulated expression. Here we show for the first time that loss of cell attachment can rapidly drive *RHOV* de novo transcription in tumor cells. These findings correlate with studies investigating the role of RHOV during neural crest development in *xenopus*, where neural tube cells upregulated RHOV expression prior to detachment and subsequent migration and differentiation (19, 20). Interestingly, we observed that MG132 treatment led to a dramatic accumulation of recombinant RHOV protein (Fig. S9). While this requires further investigation this suggests an additional layer of post-translational regulation of RHOV. Together, these findings support the hypothesis that RHOV is rapidly induced in response to cellular detachment, but is also removed via proteasomal degradation to prevent aberrant activation, suggesting that RHOV may be a tightly regulated atypical Rho GTPase during metastatic progression. For this reason RHOV has likely evaded prior detection in ovarian cancer studies. Indeed, RHOV has surfaced as a top functional hit only in unbiased CRISPR-based functional screens, including in models of breast cancer metastasis and Zika virus entry (25, 58). Aside from those studies, RHOV’s role in cancer has been explored only through targeted approaches, primarily in lung cancer, where it was shown to promote migration and invasion (22, 23). Our work now extends these observations, placing RHOV within a mechanistically defined framework of detachment-induced stress adaptation, cytoskeletal regulation, and metastatic fitness, driven by a rare combination of constitutive activity, membrane localization, and tightly controlled expression.

Further highlighting the atypical nature of RHOV, its membrane localization is similarly distinct. Unlike most Rho GTPases, which are prenylated and regulated by GDP dissociation inhibitors (GDI), RHOV is palmitoylated at its C-terminus, rendering it resistant to GDI-mediated cytosolic sequestration and stably associated with cellular membranes upon expression (49, 59). This structural feature emphasizes the necessity of tight transcriptional and proteasomal control, as RHOV, once expressed, becomes membrane-localized and hypothesized to be functionally active by default, due to its rapid GTP hydrolysis cycle (16–18). To support this notion, we find that WT-RHOV can rescue signaling and metastatic functions in RHOV-KO cells similar to the constitutively active G40V mutant, while the ΔMEM mutant acts similar to the dominant negative rescue construct. These findings highlight two potential therapeutic vulnerabilities. First RHOV’s GTP-binding domain presents one plausible target, however, the high degree of structural homology within the GTP-binding domains across Rho family GTPases might pose significant challenges related to specificity and off-target effects. Alternatively, targeting RHOV’s unique membrane localization signal may offer a more selective therapeutic strategy. Unlike other Rho GTPases, RHOV possesses a distinct, 16-residue carboxyl-terminal extension ending with a palmitoylation motif (CFV). Targeting this unique palmitoylation site or the adjacent C-terminal sequence could therefore disrupt RHOV membrane localization and selectively impair its oncogenic function, offering higher specificity and potentially fewer off-target effects. Future studies exploring this novel therapeutic approach may yield valuable insights into targeted intervention strategies for ovarian cancer metastasis.

We also report novel functional consequences of rapid, detachment-induced RHOV expression are unique and significant to peritoneal metastasis. RHOV-deficient cells exhibit failure to resist anoikis, form structurally loose and disorganized aggregates in suspension, and are markedly impaired in their ability to clear mesothelial layers, migrate, and invade through extracellular matrices. These phenotypes span multiple key steps of the transcoelomic metastatic cascade. We now show that these mechanisms are supported by RHOV and thus initiating within hours of detachment. This functional dependence on RHOV likely explains the dramatic loss of metastatic burden observed in our intraperitoneal (IP) xenograft model. In this setting, where cells are injected directly into the peritoneal space, the absence of RHOV renders cells susceptible to anoikis and inability to form MCAs In turn this limits their capacity to engage with and clear the mesothelium lining the peritoneal cavity, culminating in minimal tumor colonization of the peritoneum and omentum. Similarly, in the orthotopic intra-bursal model, RHOV knockout resulted in smaller primary tumors and a complete absence of omental and peritoneal metastases. While the reduced tumor size may partially reflect modestly impaired proliferation or reduced local invasion into adjacent stromal tissue, the absence of secondary lesions strongly supports a failure of dissemination and/or colonization. These findings further reinforce RHOV’s essential role in enabling cells to survive and adapt during the earliest stages of metastatic escape.

Mechanistically, RHOV coordinates cytoskeletal remodeling and downstream signaling during metastatic adaptation. RHOV engages the PAK–c-Jun signaling axis, driving transcriptional programs that support motility and cytoskeletal machinery. While prior studies have implicated RHOV in cytoskeletal regulation and filopodia formation in breast cancer cells (25), our findings represent the first demonstration of this pathway being functionally engaged in ovarian cancer and in response to detachment. RHOV-mediated actin cytoskeletal remodeling likely exerts broader functional consequences on cellular structural integrity, locomotion and contractility, all of which are essential for cell survival following detachment, MCA formation and invasive efficiency of metastasizing cancer cells. Alterations in actin architecture can directly influence the localization, clustering, and signaling efficacy of adhesion molecules, thereby modulating their capacity to transduce essential survival signals during a-i. Indeed, our RNA-sequencing analysis revealed significant disruption of cell-cell adhesion and junction-related transcriptional pathways in RHOV-deficient cells specifically under anchorage-independent conditions. This finding correlates strongly with the observed phenotypes, including looser spheroid formation, impaired aggregation, and reduced binding efficiency to collagen-rich mesothelial substrates. Furthermore, these observations resonate with previous studies implicating RHOV in the regulation of E-cadherin dynamics during Xenopus epiboly (21), underscoring a conserved role for RHOV in mediating adhesion-related signaling events.

Collectively, this work sheds new light on RHOV, an understudied and atypical member of the Rho GTPase family, and positions it as a critical mediator of metastatic fitness in ovarian cancer. These findings further underscore a broader biological principle: that tumor cells selectively deploy distinct Rho GTPases to navigate specific microenvironmental challenges, with RHOV emerging as the first identified detachment-sensitive Rho GTPase. While multiple GTPases have been implicated at different stages of cancer progression in both adherent and anchorage-independent states (14), our study is the first to identify RHOV as a detachment sensitive Rho GTPase rapidly engaged in early anchorage independent conditions. We propose that its constitutive activity and tight regulation enables rapid morphological and transcriptional responses that position cancer cells to adapt to stress associated with detachment and initiate pro-metastatic signaling, necessary for peritoneal dissemination of ovarian cancer cells. Finally, our study highlights that early transcriptional adaptations, immediately following matrix detachment, have lasting consequences for metastatic progression. This recognition calls for a shift in how we conceptualize and study metastatic adaptations by highlighting the discovery that immediate cellular responses to detachment prime tumor cells for successful dissemination. Ultimately, these insights open the door for better understanding the biological bases of the earliest, and perhaps most vulnerable, stages of metastatic spread.

## Materials & Methods

### Patient Ascites-Derived Cells

Primary ascites-derived cells were isolated from malignant ascites obtained from patients with high-grade serous ovarian cancer treated at Magee-Womens Hospital of UPMC. Specimen acquisition was approved by the University of Pittsburgh Institutional Review Board (IRB) (Protocol #STUDY19060197) and facilitated through the ProMark Biospecimen Bank at Magee-Womens Research Institute. Informed consent was obtained by the clinical care teams prior to specimen collection, and all samples were de-identified before laboratory processing. Ascites samples were processed immediately upon receipt for ascites-derived epithelial cell isolation and cultured as previously described (60). Cells were maintained at 37 °C in a humidified incubator with 5% CO₂ in MCDB/M199 medium supplemented with 10% fetal bovine serum (FBS) and penicillin-streptomycin.

### Patient-Matched Tumor Specimens

Archived, patient-matched tissue specimens, including normal fallopian tube or ovary, primary ovarian tumor, and omental metastases, from individuals diagnosed with high-grade serous ovarian cancer were obtained via an honest broker from the ProMark Biospecimen Bank at the University of Pittsburgh Magee-Womens Research Institute. The study protocol was approved by the University of Pittsburgh Institutional Review Board (IRB), (Protocol # STUDY19060197). All specimen collection and use adhered to ethical guidelines established by the World Medical Association’s Declaration of Helsinki and the U.S. Department of Health and Human Services Belmont Report.

### Cell Lines and Culture Conditions

OVCA433 cells were kindly provided by Dr. Susan K. Murphy (Duke University) and cultured in RPMI 1640 medium (Corning™, 10-040-CV) supplemented with 10% fetal bovine serum (FBS). SKOV3 cells stably expressing luciferase were a gift from Dr. Mythreye Karthikeyan (University of Alabama at Birmingham) and maintained in RPMI 1640 medium (Corning™, 10-040-CV) with 10% FBS. OVCAR3 cells expressing luciferase were a gift from Dr. Lan Coffman (University of Pittsburgh) and cultured in RPMI 1640 (Gibco™, 11875) supplemented with 10% FBS and 0.01 mg/mL bovine insulin (MilliporeSigma™, I0516). FT282 cells were provided by Dr. Ronny Drapkin (University of Pennsylvania) and grown in a 1:1 mixture of DMEM and Ham’s F-12 (Corning™, 10-090-CV) with 2% FBS. ZTGFP mesothelial cells, stably expressing GFP, were provided by Dr. Ioannis Zervantonakis (University of Pittsburgh) and maintained in a 1:1 mixture of Medium 199 (Corning™, 10-060-CV) and MCDB105 (Cell Applications, 117-500), supplemented with 10% FBS and 1% Penicillin-Streptomycin (Gibco™). TOV21G, ES-2, and OV90 ovarian cancer cells were purchased from the American Type Culture Collection (ATCC). TOV21G was maintained in a 1:1 mixture of MCDB 105 (Cell Applications, 117-500) and Medium 199 (Corning™, 10-060-CV) and supplemented with 15% FBS. ES-2 was cultured in McCoy’s 5A Medium (Corning, 10-050-CV) supplemented with 10% FBS. OV90 cells were maintained in a 1:1 mixture of MCDB 105 (Cell Applications, 117-500) with 1.5 g/L sodium bicarbonate (Gibco™, 25080094) and Medium 199 (Corning™, 10-060-CV) with 2.2 g/L sodium bicarbonate, supplemented with 15% FBS. Pt412 patient-derived cells were a kind gift from Dr. Ronald Buckanovich (University of Pittsburgh) and maintained in RPMI 1640 medium (Corning™, 10-040-CV) with 10% FBS. All cell lines were maintained at 37 °C in a humidified incubator with 5% CO₂, were regularly tested for mycoplasma contamination using the EZ-PCR Mycoplasma Detection Kit (Captivate Bio™, 20-700-20) and authenticated by short tandem repeat (STR) profiling (Labcorp).

### Cell Culture in Ultra-Low Attachment (ULA) Conditions

Cells were trypsinized, counted, and seeded at equal viable densities into Ultra-Low Attachment (ULA) plates (Corning) to induce anchorage-independent (a-i) growth. For RNA and protein extraction, 2 × 10⁵ cells were seeded in 2 mL of medium per well in 6-well ULA plates (Corning, 3471) and incubated for either 2 hrs (early a-i, single-cell state) or 24 hrs (later a-i, multicellular aggregate formation). Parallel control cultures representing the attached state were seeded at the same density in standard tissue culture-treated 6-well plates. For functional assays, cells were seeded at 1,000 cells per well in 96-well round-bottom ULA plates (Corning, 7007) to promote multicellular aggregate formation, or at 10,000 cells per well in 96-well flat-bottom ULA plates (Corning, 3474) in serum-free medium to maintain cells in a single-cell state. To assess the impact of dissociation method on RHOV expression, cells were enzymatically or non-enzymatically dissociated using one of the following: 0.25% Trypsin-EDTA (Fisher Scientific™, 25-200-056), TrypLE™ Express Enzyme (Thermo Scientific™, 12605028), Accutase™ (BD Biosciences, 561527), or 10 mM EDTA in PBS (Fisher Scientific™, AM9260G). Following dissociation, cells were counted and seeded at a density of 2 × 10⁵ viable cells per well in 6-well Ultra-Low Attachment (ULA) plates (Corning, 3471) in complete medium. Cells were incubated under anchorage-independent conditions for 2 hrs prior to RNA extraction for analysis of RHOV expression.

### CRISPR-Cas9 Mediated Knockout of RHOV

Isogenic RHOV-knockout (KO) derivatives were generated from the three ascites-derived ovarian cancer cell lines used in our initial screen: OVCAR3, OVCA433, and SKOV3. A multiplex CRISPR–Cas9 strategy was employed (61) to target exon 1 of RHOV, including the translation start site. Two single-guide RNA (sgRNA) were used: gRNA1 (sequence: 5’-ACAAACCCGCTGGTAGCGCT-3’) and gRNA2 (sequence: 5’-ATGGGTACCCCGCGCGCTAC-3’), each expressed from U6 promoters on separate plasmids (Addgene), each co-expressing wild-type SpCas9 (Fig. S4A). To facilitate sorting, one plasmid contained a GFP marker and the other an RFP marker. Cells were transiently co-transfected with both constructs, and dual-positive cells were identified by fluorescence microscopy and sorted as single cells into 96-well plates 48 hrs post-transfection. Co-expression of Cas9 and the two sgRNAs resulted in excision of the intervening genomic region via non-homologous end joining (NHEJ) repair (see schematic in Fig. S3A). Clonal populations were screened by genomic PCR using primers flanking the targeted locus. The outer primers used were: OUT-forward (5′-GGCCGCCTGAAACATGTAGA-3′) and OUT-reverse (5′-GCTGTGTCCCAGAGCTCAAT-3′). Successful deletion of the 296 bp region spanning exon 1 resulted in a 329 bp product, while wild-type alleles yielded a 625 bp amplicon. An additional screening PCR was performed using an internal primer (IN-forward: 5′-CAAGAGCAGCCTCATCGTCA-3′) paired with OUT-reverse; this yielded a 321 bp product in WT cells, while KO clones lacked amplification, confirming complete excision.

### siRNA-Mediated Knockdown of RHOV

For transient RHOV knockdown, cells were transfected with either a non-targeting siRNA SMARTpool control (Dharmacon™, D-001810-10-05) or a SMARTpool of siRNAs targeting RHOV (Dharmacon™, L-006374-00-0005). Transfections were performed using Lipofectamine™ RNAiMAX (Invitrogen™) according to the manufacturer’s protocol.

### Cloning and Lentiviral Transduction

Wild-type (WT) human RHOV (RefSeq: NM_133639) was subcloned from the pCMV6-Entry vector (Origene, RC211121) into the pLX304 lentiviral expression vector (Addgene, #25890) using Gateway® recombination cloning (Thermo Scientific). Both untagged and N-terminal MYC-DDK-tagged RHOV constructs were generated to assess endogenous-like behavior vs tagged RHOV constructs, respectively. Site-directed mutagenesis was used to introduce specific point mutations (G40V, S45N, and C234S) into the pLX304-RHOV WT plasmid using the Phusion Site-Directed Mutagenesis Kit (Thermo Scientific™, F541). For lentivirus production, 293FT cells were co-transfected with the pLX304-based constructs and packaging plasmids psPAX2 (Addgene, #12260) and pMD2.G (Addgene, #12259) using lipid-based transfection method. Viral supernatants were collected and used to transduce target cells (OVCA433 or SKOV3) seeded in 6-well plates the day prior. For transduction, 200 µL of viral supernatant was added per well in the presence of 0.8 µg/mL polybrene (MilliporeSigma, TR-1003) and incubated for 24 hrs. Three days post-transduction, cells were subjected to antibiotic selection using 15 µg/mL blasticidin (Thermo Scientific™, A1113903).

### RNA Sequencing and Transcriptomic Analysis

RNA sequencing was performed to identify both detachment-sensitive gene signatures and RHOV-regulated transcriptional pathways. For the detachment-response dataset, OVCAR3, OVCA433, and SKOV3 cells were cultured under standard adherent conditions or in ultra-low attachment (ULA) plates for 2 or 24 hrs to represent early and late anchorage-independent (a-i) states, respectively, with four biological replicates per condition. In a separate experiment, OVCA433 wild-type (WT) and RHOV-knockout (KO) cells were cultured under attached, early a-i (2 hrs), or late a-i (24 hrs) conditions, with three biological replicates per group. Total RNA was extracted using the Direct-zol™ RNA Miniprep Kit (Zymo Research, R2052). cDNA library preparation, sequencing, and bioinformatics analysis were performed by Novogene. Libraries were sequenced on the Illumina NovaSeq 6000 platform, and reads were aligned to the human genome using HISAT2. Differential expression analysis was conducted using the DESeq2 R package, with genes considered differentially expressed at an adjusted p-value ≤ 0.05. Functional enrichment analyses, including Gene Set Enrichment Analysis (GSEA), Gene Ontology (GO), and transcription factor analysis (ChEA), were performed using the BioJupies platform. Heat maps and volcano plots were generated in GraphPad Prism

### RNA Isolation and Semi-Quantitative Real Time RT-PCR

Total RNA was isolated using the Direct-zol™ RNA Miniprep Kit (Zymo Research, R2052) according to the manufacturer’s instructions, and cDNA was synthesized using the qScript™ cDNA Synthesis Kit (Quantabio, 95047). Semi-quantitative real-time RT-PCR was performed on a CFX Opus 96 Real-Time PCR System (Bio-Rad) using iTaq™ Universal SYBR® Green Supermix (Bio-Rad). Gene expression was normalized to the geometric mean of housekeeping genes and analyzed using the ΔΔCt method. Primer sequences for housekeeping genes included GAPDH (forward 5′-GAGTCAACGGATTTGGTCGT-3′, reverse 5′-TTGATTTTGGAGGGATCTCG-3′), ACTB (forward 5′-AGAGCTACGAGCTGCCTGAC-3′, reverse 5′-AGCACTGTGTTGGCGTACAG-3′), TBP (forward 5′-TTGGGTTTTCCAGCTAAGTTCT-3′, reverse 5′-CCAGGAAATAACTCTGGCTCA-3′), and HPRT1 (forward 5′-TGACCTTGATTTATTTTGCATACC-3′, reverse 5′-CGAGCAAGACGTTCAGTCCT-3′). Two RHOV primer sets were used: one for general RHOV detection (forward 5′-GCATTGAGCTCTGGGACACA-3′, reverse 5′-TGGTCCAGCTGAATTAGTACG-3′), and a second set optimized for detecting RHOV loss in knockout samples (forward 5′-CCTCATCGTCAGCTACACCTG-3′, reverse 5′-GAACGAAGTCGGTCAAAATCCT-3′). Primers for early anchorage-independent (a-i) response genes included FOSB (forward 5′-CAGTCCTGTGTGAGGATTAAGG-3′, reverse 5′-CACTTCTGCCCAGAGTAAAGAG-3′), EGR3 (forward 5′-GCGACCTCTACTCAGAGCC-3′, reverse 5′-CTTGGCCGATTGGTAATCCTG-3′), IER5 (forward 5′-TTTCTCGGGACTCCTACGGAA-3′, reverse 5′-GCTCCAGGGGTTCATGTCTC-3′), and NRARP (forward 5′-TCAACGTGAACTCGTTCGGG-3′, reverse 5′-ACTTCGCCTTGGTGATGAGAT-3′).

### Polysome Profiling

Polysome profiling was performed using cell lysates prepared as previously described in our prior publication (62). Briefly, cells cultured under adherent or anchorage-independent (ULA) conditions were treated with cycloheximide (100 µg/mL) for 10 minutes at 37 °C to arrest translation. After washing with ice-cold PBS containing cycloheximide, cells were lysed in polysome extraction buffer and clarified by centrifugation. The resulting supernatants were layered onto linear 20–47% sucrose gradients and subjected to ultracentrifugation at 34,000 rpm for 4 hrs and 15 minutes at 4 °C using an SW41 rotor (Beckman Coulter). Gradients were fractionated, and RNA from each fraction was extracted using TRIzol™ and analyzed by qRT-PCR following cDNA synthesis. For this study, previously generated and validated polysome fractionation samples were reused for downstream analysis.

### Tumor Xenografts

All animal procedures were approved by the Institutional Animal Care and Use Committee (IACUC) of the University of Pittsburgh and conducted in compliance with institutional guidelines (approved protocol IS00023174). Mice were housed in pathogen-free barrier facilities maintained for immunodeficient strains. For intraperitoneal (IP) xenograft experiments, SKOV3-luciferase wild-type (WT) and RHOV-knockout (RHOV-KO) cells were harvested by trypsinization, washed, and counted. A total of 1 × 10⁶ viable cells suspended in 150 µL PBS were injected intraperitoneally into female NOD scid gamma (NSG) mice (Jackson Laboratory). Mice were randomized into groups (n = 10 per group) based on pilot studies. Tumor progression was monitored every 2–3 days via bioluminescent imaging using an IVIS imaging system. Mice were injected intraperitoneally with D-Luciferin (15 mg/mL, PerkinElmer, 122799) at 10 µL/g body weight, and images were acquired 10 minutes post-injection. On day 35, mice were euthanized via CO₂ asphyxiation followed by cervical dislocation. At necropsy, ascites fluid was collected and quantified, and omentum and other visible peritoneal tumor sites were excised and weighed. Omentum was imaged *ex vivo* following additional luciferin administration. Tissues were fixed in 10% neutral-buffered formalin (Fisher Scientific, 5735), processed for paraffin embedding, sectioned, and stained with hematoxylin and eosin (H&E). For orthotopic intrabursal xenografts, OVCAR3-luciferase WT and RHOV-KO cells (1 × 10⁶) were resuspended in a mixture of 5 µL Matrigel (Corning, 354248) and 15 µL serum-free media, and injected into the ovarian bursa of NSG female mice (n = 5 per group). Mice were imaged every 3–4 days using IVIS, following the same luciferin injection and imaging protocol as described above. The study was terminated on day 57, at which point mice were euthanized and necropsied. Bioluminescence in the ovary and omentum was recorded *ex vivo*, and the number of peritoneal tumor nodules was documented.

### Western Blotting

Cells were collected either by scraping (for attached conditions) or centrifugation (for anchorage-independent conditions) and lysed in RIPA buffer (Thermo Scientific™, 89901) supplemented with protease and phosphatase inhibitor cocktails (Thermo Scientific™, 78443). Lysates were rotated at 4 °C for 30 minutes, followed by centrifugation at 21,000 × g for 30 minutes at 4 °C to clear debris. Protein concentrations were determined using the Pierce™ BCA Protein Assay Kit (Thermo Scientific™, 23225). Equal amounts of protein were resolved on SDS-PAGE gels (Bio-Rad) and transferred to nitrocellulose membranes (Fisher Scientific™, 45-004-001). Membranes were blocked for 1 hour at room temperature in 5% BSA (Fisher Scientific™, BP1600-100) prepared in TBS containing 0.1% Tween-20 (MilliporeSigma, 9005-64-5), then incubated overnight at 4 °C with primary antibodies diluted in blocking buffer. Primary antibodies included: RHOV (Proteintech, 26620-1-AP, 1:300; and Santa Cruz Biotechnology, sc-515072, 1:100), phospho-c-Jun (S63) (Cell Signaling Technology, 2361, 1:1000), total c-Jun (Cell Signaling Technology, 9165, 1:1000), phospho-PAK1+PAK2+PAK3 (phospho S144 + S154 + S144 + S141) (Abcam, ab40795, 1:1000), total PAK1 (Invitrogen, 71-9300, 1:1000), total PAK2 (Cell Signaling Technology, 2608, 1:1000), phospho-PAK4 (Ser474)/PAK5 (Ser602)/PAK6 (Ser560) (Cell Signaling Technology, 3241, 1:1000), total PAK4 (Cell Signaling Technology, 62690, 1:1000), total PAK6 (Thermo Scientific, PA5-34938, 1:1000), FLAG (Sigma-Aldrich, F3165, 1:1000), Vinculin (Sigma-Aldrich, V9131, 1:1000). After washing with TBST (0.1% Tween-20 in TBS), membranes were incubated with either HRP-conjugated secondary antibodies (Cytiva, NA931 and NA934, 1:10,000) for chemiluminescent detection or fluorescently labeled secondary antibodies (LI-COR Biosciences, 926-32211 and 926-68072, 1:10,000) for near-infrared imaging. Chemiluminescent signals were developed using SuperSignal™ West Femto Maximum Sensitivity Substrate (Thermo Scientific™, 34096) and visualized with the ChemiDoc XRS+ system (Bio-Rad). For fluorescence detection, membranes were imaged using the Odyssey CLx imaging system (LI-COR).

### Actinomycin D Treatment

For mRNA stability analysis, transcription was inhibited using actinomycin D (Sigma-Aldrich) at a final concentration of 10 μg/mL. Cells were pre-treated with actinomycin D for 30 minutes under attached conditions before being transferred to ultra-low attachment plates, where they were maintained in a-i in the continued presence of the inhibitor for subsequent time-point collection.

### Proteasomal Inhibition

Cells were treated with 10 µM MG132 (Sigma-Aldrich, 474790; Calbiochem) or an equivalent volume of DMSO as a vehicle control overnight under standard culture conditions. After treatment, cells were harvested and lysed, and protein extracts were analyzed by Western blotting to evaluate the effects of proteasomal inhibition on target protein levels.

### Cellular Proliferation

Cell proliferation assays were performed using the FluoReporter™ Blue Fluorometric dsDNA Quantitation Kit (Invitrogen™, F2962) according to the manufacturer’s instructions. Briefly, OVCAR3, OVCA433, and SKOV3 cells were seeded in 96-well plates at densities of 2000, 1000, and 500 cells per well, respectively, in a volume of 200 μL of standard culture medium. Plates were harvested at 24-hour intervals over a period of three days. At each time point, culture medium was removed, and plates were stored at -80 °C. Subsequently, plates were processed with Hoechst 33258 staining for dsDNA quantification. Fluorescence intensity was measured using a Victor X fluorescence microplate reader (PerkinElmer) with excitation and emission wavelengths set at 360 nm and 460 nm, respectively. Proliferation rates were calculated by normalizing fluorescence intensity values to the initial measurement taken at day 1.

### Live/Dead Staining

Cell viability was assessed using Calcein AM and ethidium homodimer staining. For viability assays in flat-bottom conditions, cells were seeded at a density of 10,000 cells per well in 96-well flat-bottom Ultra-Low Attachment (ULA) plates (Corning, 3474) in serum-free medium to maintain single cells or small clusters. Cells were incubated for 2, 24, 48, and 72 hrs prior to staining. For round-bottom conditions, cells were seeded at a density of 1,000 cells per well in 96-well round-bottom ULA plates (Corning, 7007) in normal culture medium to form single aggregates per well and incubated for 72 hrs prior to staining. Cells were stained with 2 µM Calcein AM and 4 µM ethidium homodimer (Sigma) diluted in PBS and incubated at 37°C for 30 minutes. Images were captured immediately afterward using a Leica Thunder Imager to determine the proportion of live (Calcein-positive, green) and dead (ethidium homodimer-positive, red) cells.

### Caspase 3/7 Activity

Caspase activity was evaluated using the Caspase-Glo® 3/7 Assay (Promega, G8981) following the manufacturer’s protocol. Cells were seeded in 96-well flat-bottom ultra-low attachment (ULA) plates as described above, and caspase activity was measured after 48 hrs under anchorage-independent (a-i) conditions. An equal volume of Caspase-Glo® 3/7 reagent was added directly to each well, and plates were incubated for 30 minutes at room temperature in the dark. Luminescence was recorded using a plate reader, and background signal from medium-only wells was subtracted from all measurements.

### Clonogenic Survival

Clonogenic assays were performed to assess single-cell survival following detachment. Briefly, 100 cells per well were seeded into standard 6-well tissue culture plates and maintained under appropriate growth conditions. After colony formation (10–14 days), cells were fixed and stained with 0.05% crystal violet solution. Colonies were imaged and quantified manually.

### Spheroid Compaction

Cells were transiently labeled with CellTrace CFSE (Invitrogen™, C34554) and seeded at a density of 1,000 cells per well in 200 µL of culture medium in 96-well round-bottom Ultra-Low Attachment (ULA) plates (Corning, 7007). Spheroid formation was monitored by time-lapse imaging using the IncuCyte S3 live-cell analysis system (Sartorius), with images acquired every hour over a 24-hour period. Spheroid area was measured at each time point and normalized to the 1-hour time point, used as the baseline to account for initial cell settling within the imaging field.

### Mesothelial Clearance

Mesothelial clearance assays were performed by initially preparing mesothelial monolayers (38). Flat-bottom 96-well plates were coated with calf skin collagen type I (Sigma-Aldrich, 8919) at a concentration of 0.05 mg/mL, incubated at 37°C, and washed three times with PBS prior to cell seeding. Mesothelial cells were then trypsinized, counted, and plated at a density of 50,000 cells per well in culture medium. Monolayer formation was confirmed 24 hrs post-seeding. For ovarian cancer spheroid formation, SKOV3 wild-type (WT) and RHOV-knockout (RHOV-KO) cells were trypsinized, counted, transiently labeled with CellTrace Far Red (Invitrogen™, C34572), and seeded at a density of 1000 cells per 200 µL medium in round-bottom 96-well Ultra-Low Attachment (ULA) plates (Corning, 7007). The plates were incubated overnight to allow spheroid formation. The following day, a single spheroid was carefully transferred onto each pre-established mesothelial monolayer. Images were captured at multiple time points (0, 4, 8, 12, and 24 hrs for initial clearance, and at 120 hrs for long-term displacement) using a Leica Thunder Imager. Areas cleared by the spheroids were quantified using FIJI software (NIH).

### Wound healing

Wound healing assays were performed using the ibidi µ-Dish with Culture-Insert 3 Well (ibidi, Cat. No. 80366), which features two defined 500 µm cell-free gaps per insert to facilitate standardized migration analysis. SKOV3 cells were seeded at a density of 30,000 cells per chamber in 70 µL of complete medium and incubated for 24 hrs to allow monolayer formation. Following insert removal, wells were gently washed with PBS to eliminate residual serum, and replaced with serum-free medium. Cells were imaged every 24 hrs for 5 days to monitor wound closure, with media changed every 48 hrs.

### Transwell Invasion

Invasion assays were conducted using Boyden chambers with 8 µm pore-size Transwell inserts coated with Matrigel at 0.4 mg/mL. The coating was allowed to solidify for 3 hrs at 37 °C. For baseline invasion assays, 5 × 10⁴ were suspended in serum-free medium and seeded into the upper chamber, while the lower chamber contained complete growth medium (RPMI with 10% FBS) to act as a chemoattractant. Cells were incubated overnight to allow invasion through the Matrigel-coated membrane. After the incubation period, non-invading cells were removed from the upper side of the membrane using a cotton swab. Invaded cells on the lower surface were stained with 0.5% crystal violet solution, washed, and air-dried. Stained cells were visualized under a microscope, and invasion was quantified by counting cells in multiple representative fields.

### Immunofluorescence Staining

For staining of adherent cells, cells were seeded into 8-well chambered slides (Falcon™, 08-774-26) and allowed to attach overnight. The next day, cells were fixed with 4% paraformaldehyde (BTC BeanTown Chemical, 30525-89-4) for 10 minutes at room temperature, permeabilized with 0.25% Triton X-100 (Fisher Scientific™, BP151500) in PBS (Corning™, 21-040-CV) for 10 minutes and blocked with Pierce™ SuperBlock™ T20 (TBS) Blocking Buffer (Thermo Scientific™, 37535) for 30 minutes. Cells were then washed three times with PBS and incubated with Alexa Fluor™ 647-conjugated phalloidin (Thermo Fisher Scientific™, A22287) for 1 hour at room temperature in the dark. After washing, coverslips were mounted using ProLong™ Gold Antifade Mountant with DAPI (Cell Signaling Technology, 8961) and cured overnight in the dark at room temperature. For wound healing experiments, cells were fixed directly in their culture wells with 4% paraformaldehyde for 10 minutes, permeabilized with 0.25% Triton X-100 in PBS, and blocked with SuperBlock™ for 30 minutes at room temperature. Cells were stained with Alexa Fluor™ 647-conjugated phalloidin and DAPI for 1 hour at room temperature in the dark, followed by PBS washes. For immunofluorescence staining of early anchorage-independent (a-i) cells, single-cell suspensions grown in ultra-low attachment (ULA) conditions were collected, fixed with 4% paraformaldehyde, and permeabilized in 0.5% Triton X-100 in TBS for 30 minutes with rotation at room temperature. Cells were blocked in SuperBlock™ for 1 hour at room temperature and then washed three times with 0.1% Triton X-100 in TBS. After a 15-minute incubation with DAPI and Alexa Fluor™ 647-conjugated phalloidin, cells were washed again and mounted using fluorescence-compatible mounting medium. Images for all conditions were acquired using either a Leica Thunder Imager or Leica DMI8 confocal microscope.

### Statistical Analysis

Unless specified otherwise, data represent the mean ± standard error of the mean (SEM) from a minimum of three independent experiments. Statistical analyses were conducted using GraphPad Prism version 10 (GraphPad Software, San Diego, CA), with methods selected according to the experimental design described. A p-value of less than 0.05 was considered statistically significant.

## Acknowledgements

The authors would like to thank Weihua Pan and Ben Yankasky for their technical assistance. This work was supported by the NIH R01CA242021 (to NH), NIH R01CA230628 (to KM and NH), NIH training grant 2T32HL110849-11A1 (to SW), and Department of Defense Ovarian Cancer Research Program HT94252310207 (to IKZ). Lauren Borho and Dr. Francesmary Mudugno kindly assisted as honest brokers to access patient specimens. The ProMark tissue bank is supported by NIH SPORE P50CA272218. This project used the Hillman Cancer Center Cytometry Facility, Animal Facility and In vivo Imaging Facility that are supported in part by award P30CA047904.

## References

1. Siegel RL, Kratzer TB, Giaquinto AN, Sung H, Jemal A. Cancer statistics, 2025. CA Cancer J Clin. 2025;75(1):10–45.

2. Tan DS, Agarwal R, Kaye SB. Mechanisms of transcoelomic metastasis in ovarian cancer. Lancet Oncol. 2006;7(11):925–34.

3. Rickard BP, Conrad C, Sorrin AJ, Ruhi MK, Reader JC, Huang SA, et al. Malignant Ascites in Ovarian Cancer: Cellular, Acellular, and Biophysical Determinants of Molecular Characteristics and Therapy Response. Cancers (Basel). 2021;13(17).

4. Huang H, Li YJ, Lan CY, Huang QD, Feng YL, Huang YW, et al. Clinical significance of ascites in epithelial ovarian cancer. Neoplasma. 2013;60(5):546–52.

5. Cai Q, Yan L, Xu Y. Anoikis resistance is a critical feature of highly aggressive ovarian cancer cells. Oncogene. 2014;34(25):3315–24.

6. Uruski P, Mikuła-Pietrasik J, Pakuła M, Budkiewicz S, Drzewiecki M, Gaiday AN, et al. Malignant Ascites Promote Adhesion of Ovarian Cancer Cells to Peritoneal Mesothelium and Fibroblasts. International Journal of Molecular Sciences. 2021;22(8):4222.

7. Ford CE, Werner B, Hacker NF, Warton K. The untapped potential of ascites in ovarian cancer research and treatment. British Journal of Cancer. 2020;123(1):9–16.

8. Farsinejad S, Cattabiani T, Muranen T, Iwanicki M. Ovarian Cancer Dissemination-A Cell Biologist’s Perspective. Cancers (Basel). 2019;11(12).

9. Skubitz KMB, Matthew PB, Stefan EP, Amy PN. Disaggregation and invasion of ovarian carcinoma ascites spheroids. Journal of Translational Medicine. 2006;4(1):1–16.

10. Shield K, Ackland ML, Ahmed N, Rice GE. Multicellular spheroids in ovarian cancer metastases: Biology and pathology. Gynecol Oncol. 2009;113(1):143–8.

11. Al Habyan S, Kalos C, Szymborski J, McCaffrey L. Multicellular detachment generates metastatic spheroids during intra-abdominal dissemination in epithelial ovarian cancer. Oncogene. 2018;37(37):5127–35.

12. Aspenstrom P, Fransson A, Saras J. Rho GTPases have diverse effects on the organization of the actin filament system. Biochem J. 2004;377(Pt 2):327–37.

13. Sahai E, Marshall CJ. RHO-GTPases and cancer. Nat Rev Cancer. 2002;2(2):133–42.

14. Crosas-Molist E, Samain R, Kohlhammer L, Orgaz JL, George SL, Maiques O, et al. Rho GTPase signaling in cancer progression and dissemination. Physiol Rev. 2022;102(1):455–510.

15. Sorokina EM, Chernoff J. Rho-GTPases: new members, new pathways. J Cell Biochem. 2005;94(2):225–31.

16. Shutes A, Berzat AC, Cox AD, Der CJ. Atypical mechanism of regulation of the Wrch-1 Rho family small GTPase. Curr Biol. 2004;14(22):2052–6.

17. Hodge RG, Ridley AJ. Regulation and functions of RhoU and RhoV. Small GTPases. 2020;11(1):8–15.

18. Shutes A, Berzat AC, Chenette EJ, Cox AD, Der CJ. Biochemical analyses of the Wrch atypical Rho family GTPases. Methods Enzymol. 2006;406:11–26.

19. Guemar L, de Santa Barbara P, Vignal E, Maurel B, Fort P, Faure S. The small GTPase RhoV is an essential regulator of neural crest induction in Xenopus. Dev Biol. 2007;310(1):113–28.

20. Faure S, Fort P. Atypical RhoV and RhoU GTPases control development of the neural crest. Small GTPases. 2015;6(4):174–7.

21. Tay HG, Ng YW, Manser E. A Vertebrate-Specific Chp-PAK-PIX Pathway Maintains E-Cadherin at Adherens Junctions during Zebrafish Epiboly. PLOS ONE. 2010;5(4):e10125.

22. Zhang D, Jiang Q, Ge X, Shi Y, Ye T, Mi Y, et al. RHOV promotes lung adenocarcinoma cell growth and metastasis through JNK/c-Jun pathway. Int J Biol Sci. 2021;17(10):2622–32.

23. Chen H, Xia R, Jiang L, Zhou Y, Xu H, Peng W, et al. Overexpression of RhoV Promotes the Progression and EGFR-TKI Resistance of Lung Adenocarcinoma. Front Oncol. 2021;11:619013.

24. Shepelev MV, Korobko IV. The RHOV gene is overexpressed in human non-small cell lung cancer. Cancer Genet. 2013;206(11):393–7.

25. Jin ML, Gong Y, Ji P, Hu X, Shao ZM. In vivo CRISPR screens identify RhoV as a pro-metastasis factor of triple-negative breast cancer. Cancer Sci. 2023;114(6):2375–85.

26. Kim YS, Gupta Vallur P, Jones VM, Worley BL, Shimko S, Shin DH, et al. Context-dependent activation of SIRT3 is necessary for anchorage-independent survival and metastasis of ovarian cancer cells. Oncogene. 2020;39(8):1619–33.

27. Shonibare Z, Monavarian M, O’Connell K, Altomare D, Shelton A, Mehta S, et al. Reciprocal SOX2 regulation by SMAD1-SMAD3 is critical for anoikis resistance and metastasis in cancer. Cell Rep. 2022;40(4):111066.

28. 28. DepMap Public 25Q2 [Internet]. Broad. 2025. Available from: depmap.org.

29. Arafeh R, Shibue T, Dempster JM, Hahn WC, Vazquez F. The present and future of the Cancer Dependency Map. Nat Rev Cancer. 2025;25(1):59–73.

30. Domcke S, Sinha R, Levine DA, Sander C, Schultz N. Evaluating cell lines as tumour models by comparison of genomic profiles. Nat Commun. 2013;4:2126.

31. Guo H, Golczer G, Wittner BS, Langenbucher A, Zachariah M, Dubash TD, et al. NR4A1 regulates expression of immediate early genes, suppressing replication stress in cancer. Mol Cell. 2021;81(19):4041–58 e15.

32. Capellero S, Erriquez J, Battistini C, Porporato R, Scotto G, Borella F, et al. Ovarian Cancer Cells in Ascites Form Aggregates That Display a Hybrid Epithelial-Mesenchymal Phenotype and Allows Survival and Proliferation of Metastasizing Cells. Int J Mol Sci. 2022;23(2).

33. Goyeneche A, Lisio MA, Fu L, Srinivasan R, Valdez Capuccino J, Gao ZH, et al. The Capacity of High-Grade Serous Ovarian Cancer Cells to Form Multicellular Structures Spontaneously along Disease Progression Correlates with Their Orthotopic Tumorigenicity in Immunosuppressed Mice. Cancers (Basel). 2020;12(3).

34. Sodek KL, Ringuette MJ, Brown TJ. Compact spheroid formation by ovarian cancer cells is associated with contractile behavior and an invasive phenotype. Int J Cancer. 2009;124(9):2060–70.

35. Klymenko Y, Johnson J, Bos B, Lombard R, Campbell L, Loughran E, et al. Heterogeneous Cadherin Expression and Multicellular Aggregate Dynamics in Ovarian Cancer Dissemination. Neoplasia. 2017;19(7):549–63.

36. Iwanicki MP, Davidowitz RA, Ng MR, Besser A, Muranen T, Merritt M, et al. Ovarian cancer spheroids use myosin-generated force to clear the mesothelium. Cancer Discov. 2011;1(2):144–57.

37. Davidowitz RA, Selfors LM, Iwanicki MP, Elias KM, Karst A, Piao H, et al. Mesenchymal gene program-expressing ovarian cancer spheroids exhibit enhanced mesothelial clearance. J Clin Invest. 2014;124(6):2611–25.

38. Jazwinska DE, Cho Y, Zervantonakis IK. Enhancing PKA-dependent mesothelial barrier integrity reduces ovarian cancer transmesothelial migration via inhibition of contractility. iScience. 2024;27(6):109950.

39. Kenny HA, Nieman KM, Mitra AK, Lengyel E. The first line of intra-abdominal metastatic attack: breaching the mesothelial cell layer. Cancer Discov. 2011;1(2):100–2.

40. Kong HJ, Kwon EJ, Kwon OS, Lee H, Choi JY, Kim YJ, et al. Crosstalk between YAP and TGFbeta regulates SERPINE1 expression in mesenchymal lung cancer cells. Int J Oncol. 2021;58(1):111–21.

41. Polo-Generelo S, Rodriguez-Mateo C, Torres B, Pintor-Tortolero J, Guerrero-Martinez JA, Konig J, et al. Serpine1 mRNA confers mesenchymal characteristics to the cell and promotes CD8+ T cells exclusion from colon adenocarcinomas. Cell Death Discov. 2024;10(1):116.

42. Zeeh F, Witte D, Gadeken T, Rauch BH, Grage-Griebenow E, Leinung N, et al. Proteinase-activated receptor 2 promotes TGF-beta-dependent cell motility in pancreatic cancer cells by sustaining expression of the TGF-beta type I receptor ALK5. Oncotarget. 2016;7(27):41095–109.

43. Lin J, He Y, Chen L, Chen X, Zang S, Lin W. MYLK promotes hepatocellular carcinoma progression through regulating cytoskeleton to enhance epithelial-mesenchymal transition. Clin Exp Med. 2018;18(4):523–33.

44. Casey RC, Burleson KM, Skubitz KM, Pambuccian SE, Oegema TR, Jr., Ruff LE, et al. Beta 1-integrins regulate the formation and adhesion of ovarian carcinoma multicellular spheroids. Am J Pathol. 2001;159(6):2071–80.

45. Dhaliwal D, Shepherd TG. Molecular and cellular mechanisms controlling integrin-mediated cell adhesion and tumor progression in ovarian cancer metastasis: a review. Clin Exp Metastasis. 2022;39(2):291–301.

46. McKenzie AJ, Hicks SR, Svec KV, Naughton H, Edmunds ZL, Howe AK. The mechanical microenvironment regulates ovarian cancer cell morphology, migration, and spheroid disaggregation. Sci Rep. 2018;8(1):7228.

47. Minet E, Michel G, Mottet D, Piret JP, Barbieux A, Raes M, et al. c-JUN gene induction and AP-1 activity is regulated by a JNK-dependent pathway in hypoxic HepG2 cells. Exp Cell Res. 2001;265(1):114–24.

48. Aronheim A, Broder YC, Cohen A, Fritsch A, Belisle B, Abo A. Chp, a homologue of the GTPase Cdc42Hs, activates the JNK pathway and is implicated in reorganizing the actin cytoskeleton. Curr Biol. 1998;8(20):1125–8.

49. Chenette EJ, Abo A, Der CJ. Critical and distinct roles of amino- and carboxyl-terminal sequences in regulation of the biological activity of the Chp atypical Rho GTPase. J Biol Chem. 2005;280(14):13784–92.

50. Korobko IV, Shepelev MV. Mutations in the Effector Domain of RhoV GTPase Impair Its Binding to Pak1 Protein Kinase. Molecular Biology. 2018;52(4):598–603.

51. Shepelev MV, Korobko IV. Pak6 protein kinase is a novel effector of an atypical Rho family GTPase Chp/RhoV. Biochemistry (Mosc). 2012;77(1):26–32.

52. Weisz Hubsman M, Volinsky N, Manser E, Yablonski D, Aronheim A. Autophosphorylation-dependent degradation of Pak1, triggered by the Rho-family GTPase, Chp. Biochem J. 2007;404(3):487–97.

53. Kumar R, Sanawar R, Li X, Li F. Structure, biochemistry, and biology of PAK kinases. Gene. 2017;605:20–31.

54. Moss NM, Barbolina MV, Liu Y, Sun L, Munshi HG, Stack MS. Ovarian cancer cell detachment and multicellular aggregate formation are regulated by membrane type 1 matrix metalloproteinase: a potential role in I.p. metastatic dissemination. Cancer Res. 2009;69(17):7121–9.

55. Giannakouros P, Comamala M, Matte I, Rancourt C, Piche A. MUC16 mucin (CA125) regulates the formation of multicellular aggregates by altering beta-catenin signaling. Am J Cancer Res. 2015;5(1):219–30.

56. Kellouche S, Fernandes J, Leroy-Dudal J, Gallet O, Dutoit S, Poulain L, et al. Initial formation of IGROV1 ovarian cancer multicellular aggregates involves vitronectin. Tumor Biology. 2010;31(2):129–39.

57. Minatohara K, Akiyoshi M, Okuno H. Role of Immediate-Early Genes in Synaptic Plasticity and Neuronal Ensembles Underlying the Memory Trace. Front Mol Neurosci. 2015;8:78.

58. Luu AP, Yao Z, Ramachandran S, Azzopardi SA, Miles LA, Schneider WM, et al. A CRISPR Activation Screen Identifies an Atypical Rho GTPase That Enhances Zika Viral Entry. Viruses. 2021;13(11).

59. Chenette EJ, Mitin NY, Der CJ. Multiple sequence elements facilitate Chp Rho GTPase subcellular location, membrane association, and transforming activity. Mol Biol Cell. 2006;17(7):3108–21.

60. Shepherd TG, Theriault BL, Campbell EJ, Nachtigal MW. Primary culture of ovarian surface epithelial cells and ascites-derived ovarian cancer cells from patients. Nat Protoc. 2006;1(6):2643–9.

61. Ran FA, Hsu PD, Wright J, Agarwala V, Scott DA, Zhang F. Genome engineering using the CRISPR-Cas9 system. Nat Protoc. 2013;8(11):2281–308.

62. Kim YS, Tang PW, Welles JE, Pan W, Javed Z, Elhaw AT, et al. HuR-dependent SOD2 protein synthesis is an early adaptation to anchorage-independence. Redox Biol. 2022;53:102329.

