## Supplemental Figures for "RHOV is a Detachment-Responsive Rho GTPase Necessary for Ovarian Cancer Peritoneal Metastasis"

**Supplementary Results:**

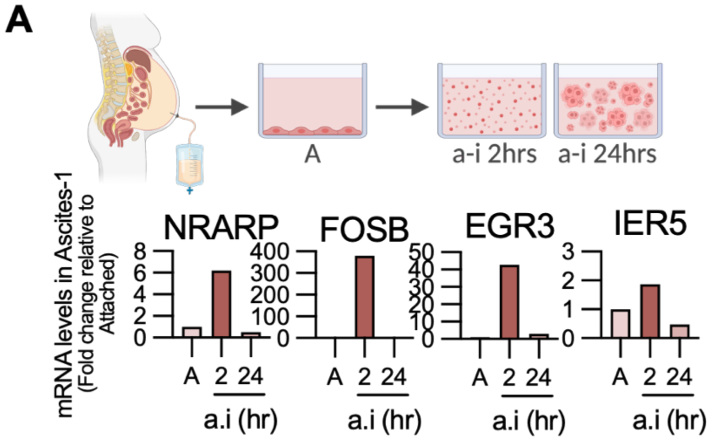


**Figure S1. Induction of immediate early response genes in early a-i in patient-derived ascites cells.**

**A,** Isolation of ascites-derived epithelial ovarian cancer cells from high grade serous ovarian cancer patient' ascites and their subsequent culture in attached vs a-i conditions. sqRT-PCR of select early response genes among the detachment-sensitive signature show upregulation in 2 hrs a-i condition in Ascites-1 patient sample.

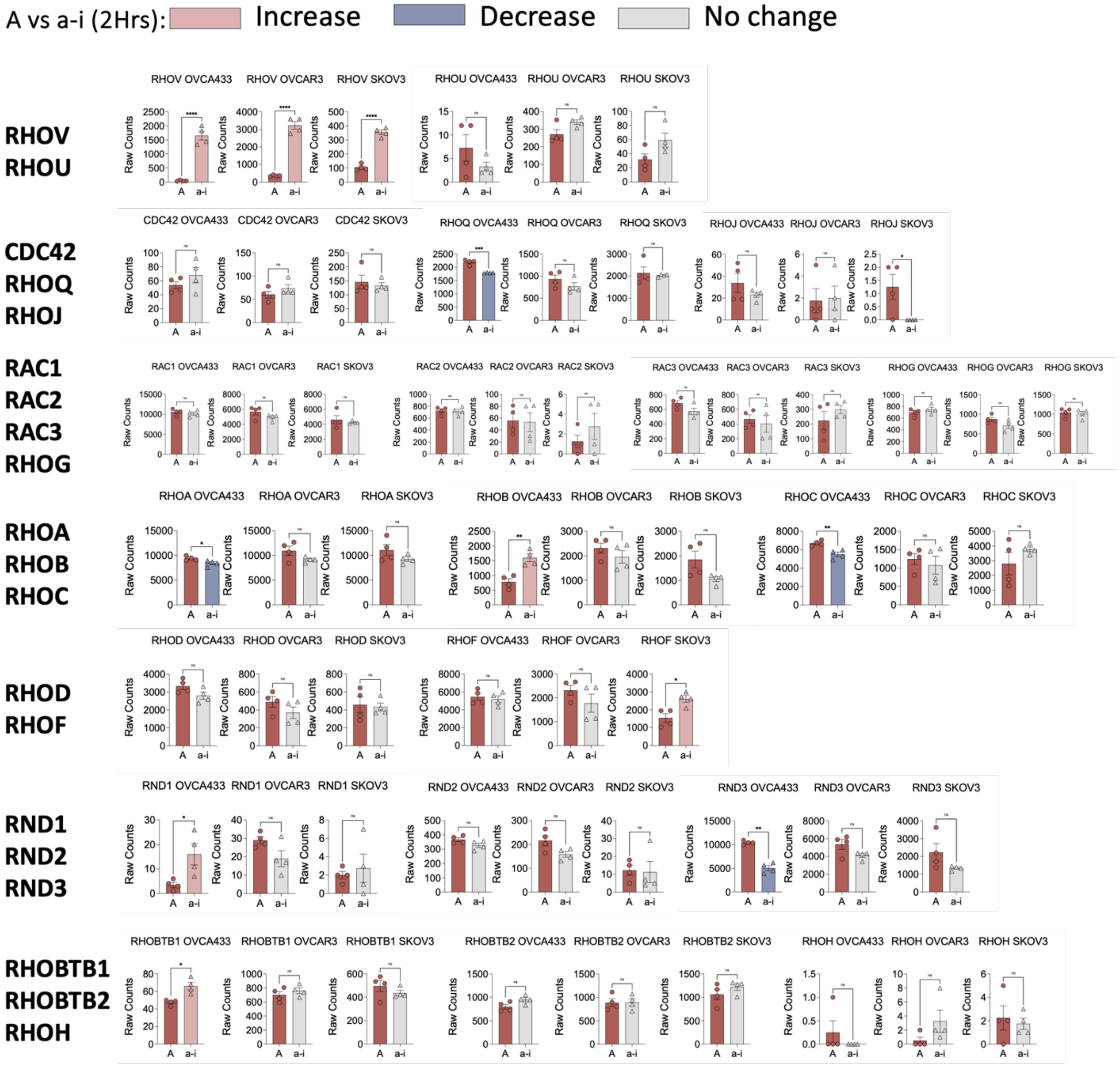


**Figure S2. RHOV is uniquely upregulated among Rho GTPases during early anchorage-independent adaptation.**

Expression levels of all 20 canonical Rho GTPases in OVCAR3, OVCA433, and SKOV3 cell lines under adherent and 2 hrs anchorage-independent conditions grouped by subfamilies. Raw counts from RNA-seq data are plotted and color coded to show significant changes across all three cell lines. RHOV is the only family member significantly and consistently upregulated in all cell lines (n=4; unpaired t-test, $^{*}P < 0.05$, $^{**}P < 0.01$, $^{***}P < 0.001$, $^{****}P<0.0001$)

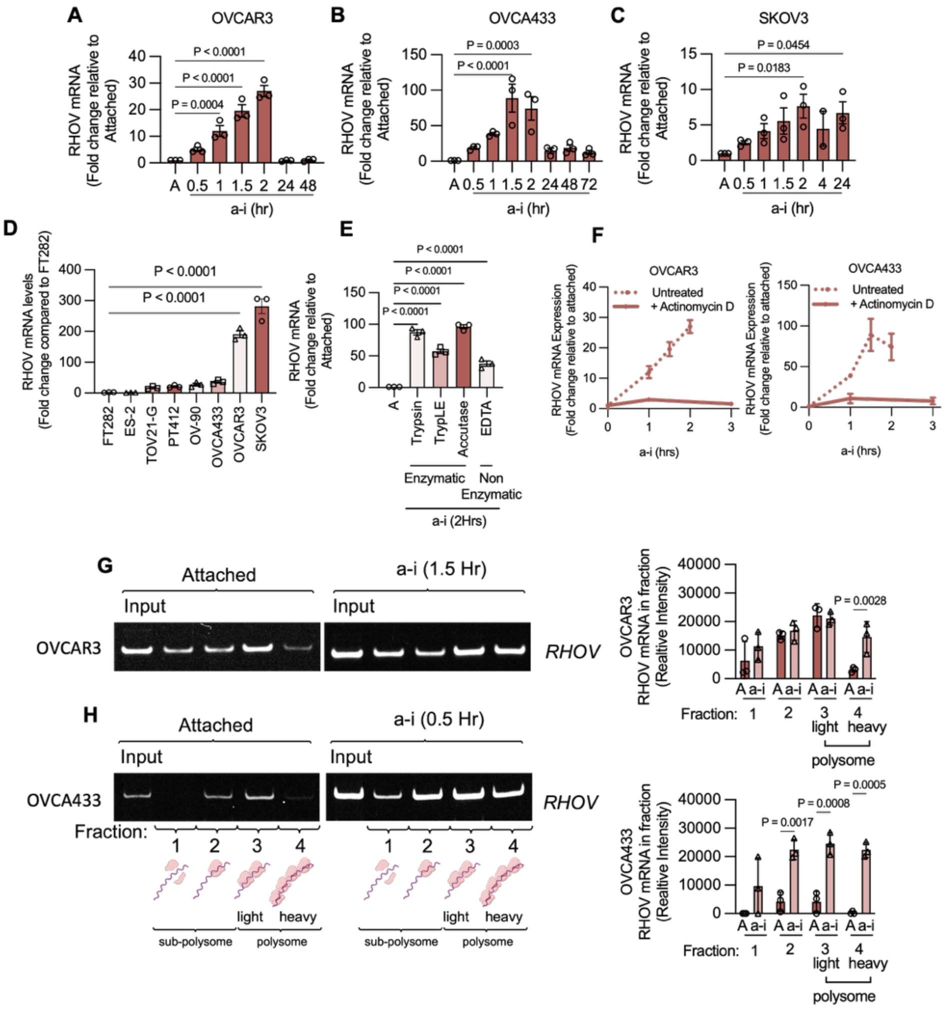


**Figure S3. Transcriptional and translational regulation of RHOV following cellular detachment.**

**A-C,** Time-course of RHOV expression post-detachment in OVCAR3 (A), OVCA433 (B), and SKOV3 (C) cells by semi quantitative RT-PCR (sqRT-PCR) shows rapid induction immediately following detachment, peaking at ~2 hrs (n=3, one way ANOVA, OVCAR3: $P < 0.0001$, OVCA433: $P < 0.0001$, SKOV3: $P < 0.05$, Dunnett post-hoc test P values are shown). **D,** sqRT-PCR showing basal RHOV expression levels in attached conditions of a panel of ovarian cancer cells compared to non-cancerous fallopian tube cell line FT282 shows enrichment in cancer cells with highest expression observed in the metastatic cell line SKOV3. (n=3; one way ANOVA $P < 0.0001$, Dunnett's post hoc test P value shown). **E,** sqRT-PCR showing RHOV induction at 2 hrs a-i following either enzymatic (trypsin, TryplE, Accutase) or non-enzymatic (EDTA) detachment in OVCA433 cells demonstrating the detachment-induced nature of RHOV upregulation independent on detachment method (n=3; one way ANOVA $P < 0.0001$, Dunnett's post hoc test P value shown).**F,** sqRT-PCR analysis of RHOV mRNA in OVCAR3 and OVCA433 cells cultured under anchorage-independent (a-i) conditions with or without pretreatment with actinomycin D (10 µg/mL, 30 min). Actinomycin D abrogates RHOV induction, indicating de novo transcription (n=3). For comparative reference, RHOV levels in untreated a-i samples are shown as a dashed line, replotted from panels (A, B). **G, H,** Polysome profiling of *RHOV* mRNA in OVCAR3 (G) and OVCA433 (H) cells at 90- and 30-minutes post-detachment, respectively. RHOV transcripts shift from light to heavy polysome fractions under a-i conditions, indicating active translation. Quantification shown for each (n=3, two way ANOVA, OVCAR3 $P=0.0042$, OVCA433 $P<0.0001$, Sidak's Post hoc test P-values shown).

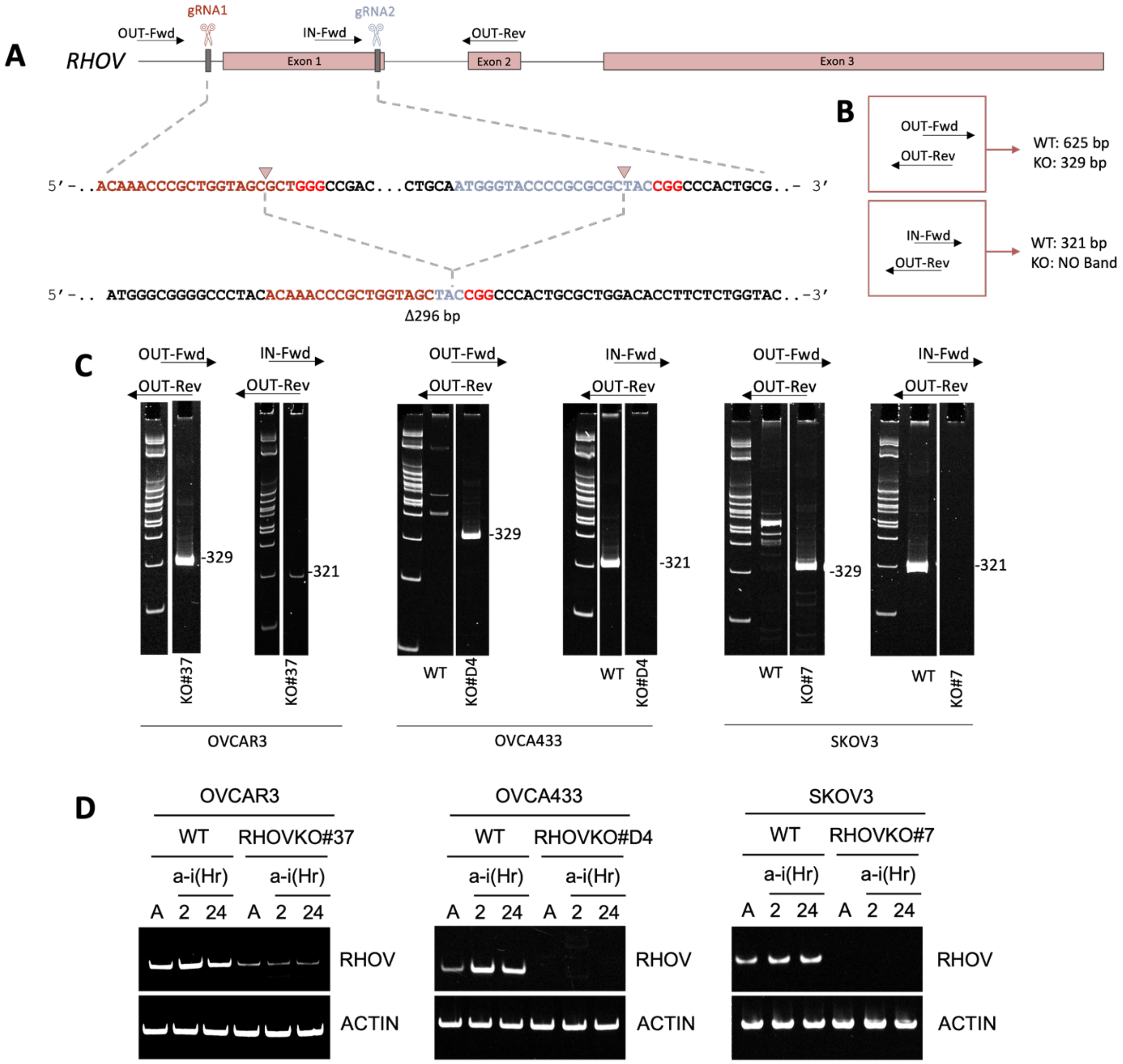


**Figure S4. CRISPR–Cas9 generation and validation of RHOV-knockout ovarian cancer cell lines.**

**A,** Schematic of dual-sgRNA CRISPR–Cas9 strategy targeting exon 1 of RHOV. Cutting sites are annotated with gRNA1/gRNA2 labeled scissors, sequence is further enlarged to show NGG sites (in red) and expected annealed product following NHEJ causing a 296bp deletion. **B,** Map representing the primer location used for subsequent RT-PCRs and expected product size for WT and KO clones. **C,** PCR screening reveals successful RHOV-KO clones in the cell lines tested. Complete deletion is observed in OVCA433 and SKOV3 clones with partial deletion in OVCAR3. **D,** RT-PCR validation of RHOV transcript loss in attached and a-i conditions for each knockout line.

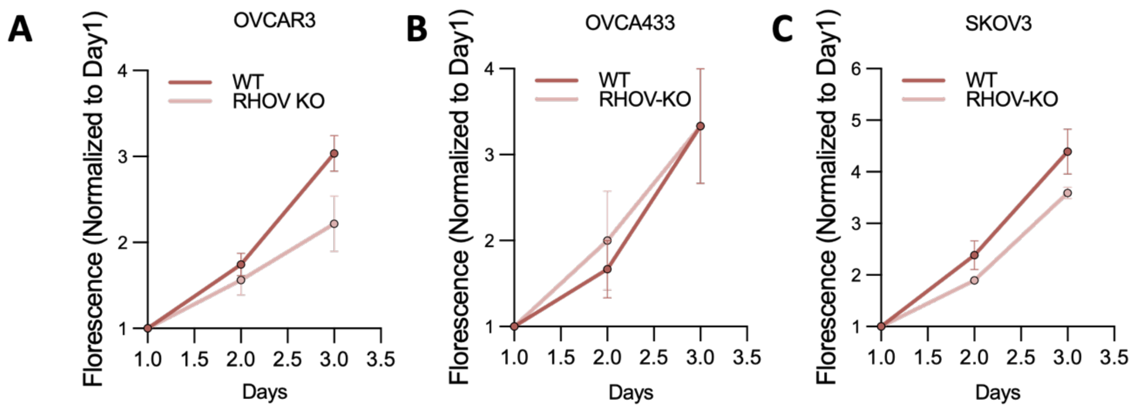


**Figure S5. Effect of RHOV deletion on proliferation under adherent conditions.**

**A-C,** Proliferation curves of WT vs RHOV-KO clones generated in all three ovarian cancer cell lines, OVCAR3 (A), OVCA433 (B), and SKOV3 (C). Cell proliferation was measured using FluoReporter dsDNA quantification and cell density relative to day 1 was plotted (n=3; two-way ANOVA, OVCAR3 P=0.04, OVCA433 P=0.86, SKOV3 P=0.1)

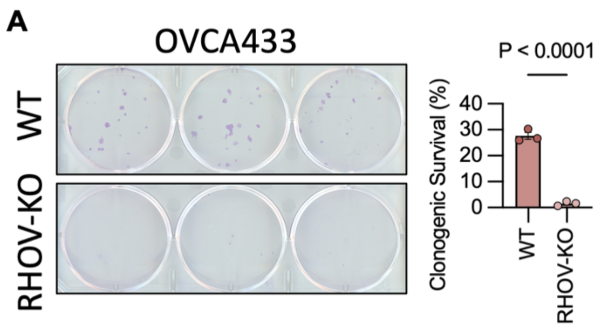


**Figure S6. The effect of RHOV-KO on OVCA433 clonogenic survival.**

**A,** Clonogenic survival assay shows reduced colony formation in RHOV-KO cells compared to WT control, demonstrating the role of RHOV in mediating single cell survival (n=3, unpaired t-test P value shown).

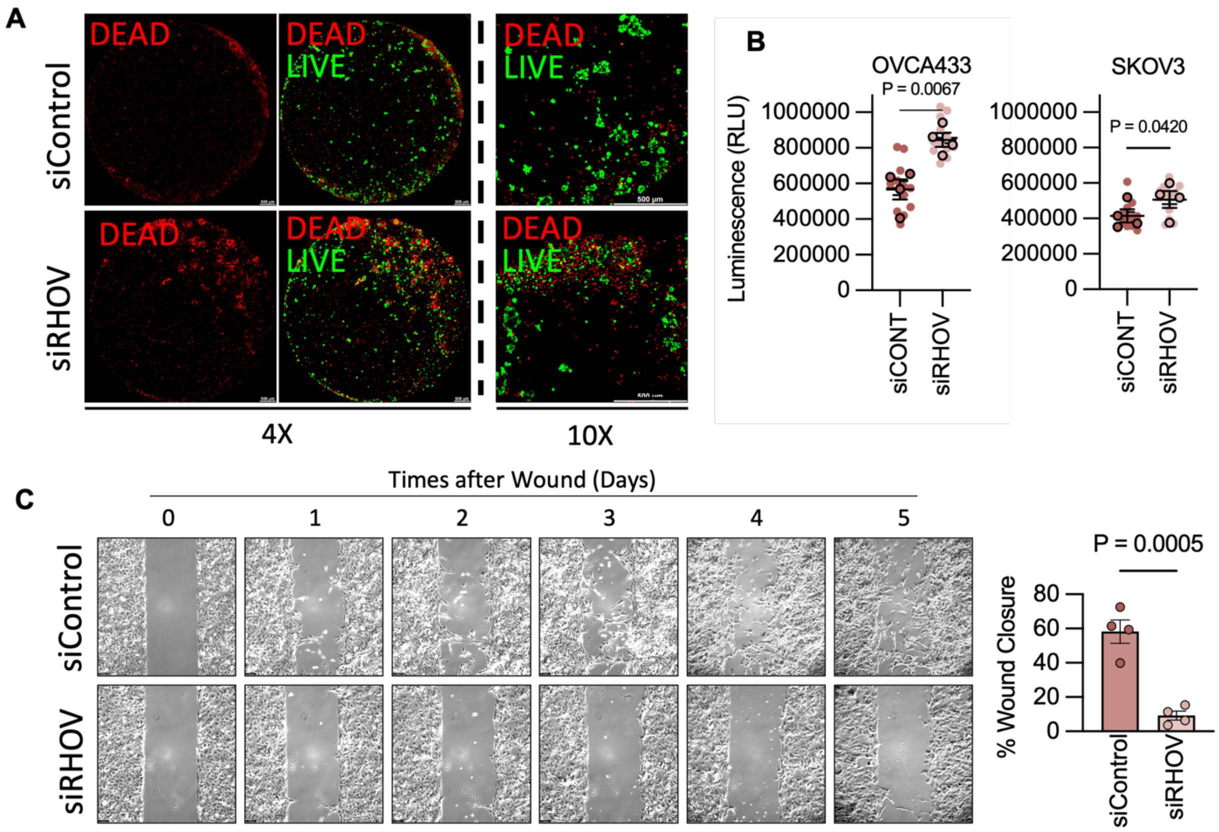


**Figure S7. siRNA-mediated RHOV depletion recapitulates RHOV-KO effects in cell culture assays.**

**A,** Representative image of increased cell death of single cells in a-i in siRHOV transfected OVCA433 cells but not in siControl following 72Hrs culture in flat bottom ultra-low attachment dishes and stained red with ethidium homodimer (dead) and green with calcein (live). Images were taken at 4x magnification and stitched (2x2 stich) to show whole well. Scale bar = 500uM. 10x magnification images are shown on left. **B,** Caspase 3/7 activity assay showing increased apoptosis in OVCA433 and SKOV3 cells transfected with siRHOV vs siControl following 72Hrs culture in flat bottom ultra-low attachment dishes further demonstrates the role of RHOV in protecting detached cells against anoikis (n=4, unpaired t-test P value shown). **C,** Representative 10x images and quantification of Wound healing assay using siRHOV in SKOV3 cells show complete abrogation of cell migration in RHOV-depleted cells, confirming that loss of RHOV impairs motility independent of CRISPR-mediated deletion. Scale bar = 100uM (n=3, unpaired t-test P value shown).

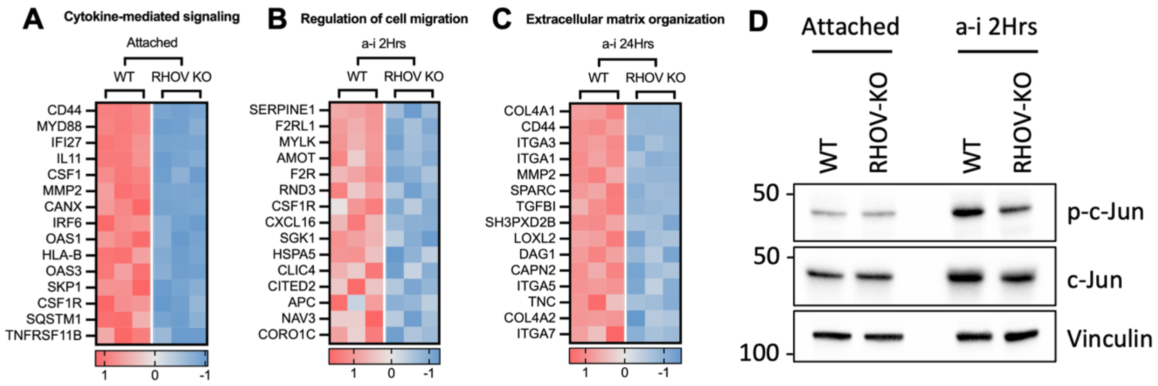


**Figure S8. Dynamic RHOV-Regulated Pathways Across Cell States and Suppression of c-Jun Signaling in RHOV-Deficient Cells**

**A,** Heat map of top downregulated genes in cytokine-mediated signaling, the top downregulated pathway in RHOV-KO cells under adherent conditions, highlighting regulation of cytokines like IFI27, IL11, CSF1, CSF1R by RHOV in attached cells. **B,** Heat map of top downregulated genes in cell migration pathway, the top downregulated pathway in RHOV-KO cells under early a-i (2Hrs) conditions, highlighting regulation of migration/metastasis related genes like MYLK, SERPINE1, F2RL1, F2R in RHOV-KO cells in early a-i. **C,** Heat map of top downregulated genes in extracellular matrix organization, the top downregulated pathway in RHOV-KO cells under late a-i (24Hrs) conditions, highlighting regulation of aggregation related genes like COL4A1, COL4A2, ITGA1, ITGA3, ITGA5, ITGA7 in RHOV-KO cells in late a-i. **D,** Representative western blot image of OVCA433 WT vs RHOV-KO cells in attached and a-i conditions showing the suppression of both total and phosphor-c-Jun in RHOV-KO cells.

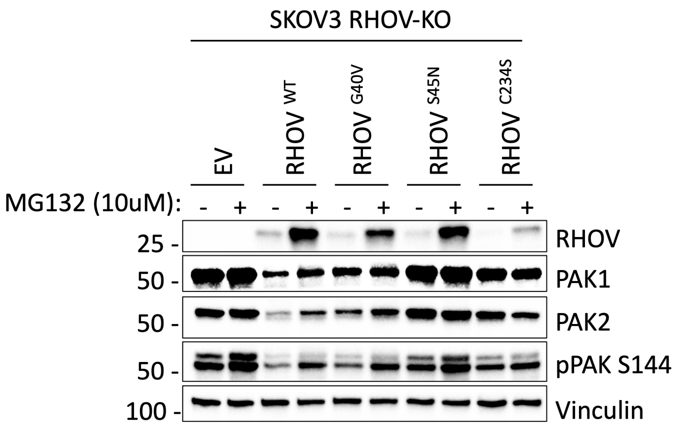


**Figure S9. Characterizing RHOV-rescue mutants following MG132 treatment.**

Western blots showing rescue of phospho- and total PAK1/2 by RHOV-WT and G40V in SKOV3-RHOV-KO cells. MG132 treatment (10uM/24Hrs) confirms phosphorylation dependent degradation of PAKs in response to RHOV expression. Blots also show noticeable accumulation of RHOV following MG132 treatment indicative of high RHOV turnover under basal conditions.
